# Trans-ethnic and ancestry-specific blood-cell genetics in 746,667 individuals from 5 global populations

**DOI:** 10.1101/2020.01.17.910497

**Authors:** Ming-Huei Chen, Laura M. Raffield, Abdou Mousas, Saori Sakaue, Jennifer E. Huffman, Tao Jiang, Parsa Akbari, Dragana Vuckovic, Erik L. Bao, Arden Moscati, Xue Zhong, Regina Manansala, Véronique Laplante, Minhui Chen, Ken Sin Lo, Huijun Qian, Caleb A. Lareau, Mélissa Beaudoin, Masato Akiyama, Traci M. Bartz, Yoav Ben-Shlomo, Andrew Beswick, Jette Bork-Jensen, Erwin P. Bottinger, Jennifer A. Brody, Frank J.A. van Rooij, Kumaraswamynaidu Chitrala, Kelly Cho, Hélène Choquet, Adolfo Correa, John Danesh, Emanuele Di Angelantonio, Niki Dimou, Jingzhong Ding, Paul Elliott, Tõnu Esko, Michele K. Evans, James S. Floyd, Linda Broer, Niels Grarup, Michael H. Guo, Andreas Greinacher, Jeff Haessler, Torben Hansen, Joanna M. M. Howson, Wei Huang, Eric Jorgenson, Tim Kacprowski, Mika Kähönen, Yoichiro Kamatani, Masahiro Kanai, Savita Karthikeyan, Fotis Koskeridis, Leslie A. Lange, Terho Lehtimäki, Markus M. Lerch, Allan Linneberg, Yongmei Liu, Leo-Pekka Lyytikäinen, Ani Manichaikul, Koichi Matsuda, Karen L. Mohlke, Nina Mononen, Yoshinori Murakami, Girish N. Nadkarni, Matthias Nauck, Kjell Nikus, Willem H. Ouwehand, Nathan Pankratz, Oluf Pedersen, Michael Preuss, Bruce M. Psaty, Olli T. Raitakari, David J. Roberts, Stephen S. Rich, Benjamin A.T. Rodriguez, Jonathan D. Rosen, Jerome I. Rotter, Petra Schubert, Cassandra N. Spracklen, Praveen Surendran, Hua Tang, Jean-Claude Tardif, Mohsen Ghanbari, Uwe Völker, Henry Völzke, Nicholas A. Watkins, Alan B. Zonderman, VA Million Veteran Program, Peter W.F. Wilson, Yun Li, Adam S. Butterworth, Jean-François Gauchat, Charleston W.K. Chiang, Bingshan Li, Ruth J.F. Loos, William J. Astle, Evangelos Evangelou, Vijay G. Sankaran, Yukinori Okada, Nicole Soranzo, Andrew D. Johnson, Alexander P. Reiner, Paul L. Auer, Guillaume Lettre

**Affiliations:** The Framingham Heart Study, National Heart, Lung and Blood Institute, Framingham, MA, USA; Population Sciences Branch, Division of Intramural Research, National Heart, Lung and Blood Institute, Framingham, MA, USA; Department of Genetics, University of North Carolina, Chapel Hill, NC, USA; Montreal Heart Institute, Montreal, Quebec, Canada; Department of Statistical Genetics, Osaka University Graduate School of Medicine, Suita, Osaka, Japan; Laboratory for Statistical Analysis, RIKEN Center for Integrative Medical Sciences, Yokohama, Kanagawa, Japan; Center for Population Genomics, Massachusetts Veterans Epidemiology Research and Information Center (MAVERIC), VA Boston Healthcare System, Boston, MA, USA; Department of Public Health and Primary Care, University of Cambridge, Cambridge, UK; MRC/BHF Cardiovascular Epidemiology Unit, Department of Public Health and Primary Care, University of Cambridge, Cambridge, UK; MRC Biostatistics Unit, University of Cambridge, Cambridge, UK; Human Genetics, Wellcome Sanger Institute, Hinxton, UK; NIHR Blood and Transplant Research Unit in Donor Health and Genomics, Strangeways Laboratory, Cambridge, UK; Division of Hematology/Oncology, Boston Children’s Hospital, Boston, MA, USA; Broad Institute of Harvard and MIT, Cambridge, MA, USA; Icahn School of Medicine at Mount Sinai, The Charles Bronfman Institute for Personalized Medicine, New York, NY, USA; Department of Medicine, Division of Genetic Medicine, Vanderbilt University, Nashville, TN, USA; Zilber School of Public Health, University of Wisconsin-Milwaukee, Milwaukee, WI, USA; Département de Pharmacologie et Physiologie, Faculty of Medicine, Université de Montréal, Montreal, Quebec, Canada; Center for Genetic Epidemiology, Department of Preventive Medicine, Keck School of Medicine, University of Southern California, Los Angeles, CA, USA; Department of Statistics and Operation Research, University of North Carolina, Chapel Hill, NC, USA; Department of Ocular Pathology and Imaging Science, Graduate School of Medical Sciences, Kyushu University, Fukuoka, Japan; Department of Biostatistics, University of Washington, Seattle, WA, USA; Population Health Sciences, Bristol Medical School, University of Bristol, Bristol, UK; Translational Health Sciences, Musculoskeletal Research Unit, Bristol Medical School, University of Bristol, Bristol, UK; Novo Nordisk Foundation Center for Basic Metabolic Research, Faculty of Health and Medical Sciences, University of Copenhagen, Copenhagen, Denmark; Hasso-Plattner-Institut, Universität Potsdam, Postdam, Germany; Department of Medicine, University of Washington, Seattle, WA, USA; Department of Epidemiology, Erasmus University Medical Center Rotterdam, Rotterdam, The Netherlands; Laboratory of Epidemiology and Population Science, National Institute on Aging/NIH, Baltimore, MD, USA; Massachusetts Veterans Epidemiology Research and Information Center (MAVERIC), VA Boston Healthcare System, Boston, MA, USA; Department of Medicine, Division on Aging, Brigham and Women’s Hospital, Boston, MA, USA; Department of Medicine, Harvard Medical School, Boston, MA, USA; Division of Research, Kaiser Permanente Northern California, Oakland, CA, USA; Department of Medicine, University of Mississippi Medical Center, Jackson, MS, USA; Health Data Research UK Cambridge, Wellcome Genome Campus and University of Cambridge, Cambridge, UK; The National Institute for Health Research Blood and Transplant Unit (NIHR BTRU) in Donor Health and Genomics at the University of Cambridge, Cambridge, UK; Human Genetics, Wellcome Sanger Institute, Saffron Walden, UK; National Institute for Health Research Cambridge Biomedical Research Centre, Cambridge University Hospitals, Cambridge, UK; British Heart Foundation Centre of Excellence, Division of Cardiovascular Medicine, Addenbrooke’s Hospital, Cambridge, UK; Section of Nutrition and Metabolism, International Agency for Research on Cancer, Lyon, France; Department of Hygiene and Epidemiology, University of Ioannina Medical School, Ioannina, Greece; Department of Internal Medicine, Section of Gerontology and Geriatric Medicine, Wake Forest School of Medicine, Winston-Salem, NC, USA; Department of Epidemiology and Biostatistics, Imperial College London, London, UK; Imperial Biomedical Research Centre, Imperial College London and Imperial College NHS Healthcare Trust, London, UK; Medical Research Council-Public Health England Centre for Environment, Imperial College London, London, UK; UK Dementia Research Institute, Imperial College London, London, UK; Health Data research UK-London, London, UK; Department of Epidemiology, University of Washington, Seattle, WA, USA; Department of Internal Medicine, Erasmus University Medical Center Rotterdam, Rotterdam, The Netherlands; Department of Neurology, University of Pennsylvania, Philadelphia, PA, USA; Institute for Immunology and Transfusion Medicine, University Medicine Greifswald, Greifswald, Germany; Division of Public Health Sciences, Fred Hutchinson Cancer Research Center, Seattle, WA, USA; Department of Genetics, Shanghai-MOST Key Laboratory of Health and Disease Genomics, Chinese National Human Genome Center and Shanghai Industrial Technology Institute (SITI), Shanghai, China; Interfaculty Institute of Genetics and Functional Genomics, University Medicine Greifswald, Greifswald, Germany; Chair of Experimental Bioinformatics, Research Group Computational Systems Medicine, Technical University of Munich, Freising-Weihenstephan, Germany; German Center for Cardiovascular Research (DZHK), Partner Site Greifswald, Greifswald, Germany; Department of Clinical Physiology, Tampere University Hospital, Tampere, Finland; Department of Clinical Physiology, Finnish Cardiovascular Research Center - Tampere, Faculty of Medicine and Health Technology, Tampere University, Tampere, Finland; Laboratory of Complex Trait Genomics, Department of Computational Biology and Medical Sciences, Graduate School of Frontier Sciences, The University of Tokyo, Tokyo, Japan; Analytic and Translational Genetics Unit, Massachusetts General Hospital, Boston, MA, USA; Department of Medicine, University of Colorado Denver, Anschutz Medical Campus, Aurora, CO, USA; Department of Clinical Chemistry, Fimlab Laboratories, Tampere, Finland; Department of Clinical Chemistry, Finnish Cardiovascular Research Center - Tampere, Faculty of Medicine and Health Technology, Tampere University, Tampere, Finland; Department of Internal Medicine, University Medicine Greifswald, Greifswald, Germany; Center for Clinical Research and Prevention, Bispebjerg and Frederiksberg Hospital, Frederiksberg, Denmark; Department of Clinical Medicine, Faculty of Health and Medical Sciences, University of Copenhagen, Copenhagen, Denmark; Department of Medicine, Division of Cardiology, Duke Molecular Physiology Institute, Duke University Medical Center, Durham, NC, USA; Center for Public Health Genomics, University of Virginia, Charlottesville, VA, USA; Department of Computational Biology and Medical Sciences, Graduate school of Frontier Sciences, The University of Tokyo, Tokyo, Japan; Division of Molecular Pathology, the Institute of Medical Sciences, The University of Tokyo, Tokyo, Japan; Institue of Clinical Chemistry and Laboratory Medicine, University Medicine Greifswald, Greifswald, Germany; Department of Cardiology, Heart Center, Tampere University Hospital, Tampere, Finland; Department of Cardiology, Finnish Cardiovascular Research Center - Tampere, Faculty of Medicine and Health Technology, Tampere University, Tampere, Finland; Department of Hematology, University of Cambridge, Cambridge Biomedical Campus, Cambridge, UK; National Health Service (NHS) Blood and Transplant, Cambridge Biomedical Campus, Cambridge, UK; Department of Laboratory Medicine and Pathology, University of Minnesota, Minneapolis, MN, USA; Departments of Epidemiology, University of Washington, Seattle, WA, USA; Department of Health Services, University of Washington, Seattle, WA, USA; Kaiser Permanente Washington Health Research Institute, Seattle, WA, USA; Centre for Population Health Research, University of Turku and Turku University Hospital, Turku, Finland; Research Centre of Applied and Preventive Cardiovascular Medicine, University of Turku, Turku, Finland; Department of Clinical Physiology and Nuclear Medicine, Turku University Hospital, Turku, Finland; Radcliffe Department of Medicine, University of Oxford, John Radcliffe Hospital, Oxford, UK; Department of Haematology, Churchill Hospital, Oxford, UK; Department of Biostatistics, University of North Carolina, Chapel Hill, NC, USA; The Institute for Translational Genomics and Population Sciences, Department of Pediatrics, The Lundquist Institute for Biomedical Innovation (formerly Los Angeles Biomedical Research Institute) at Harbor-UCLA Medical Center, Torrance, CA, USA; Department of Genetics, Stanford University School of Medicine, Stanford, CA, USA; Department of Medicine, Faculty of Medicine, Université de Montréal, Montreal, Quebec, Canada; Institute for Community Medicine, University Medicine Greifswald, Greifswald, Germany; Atlanta VA Medical Center, Decatur, GA, USA; Department of Computer Science, University of North Carolina, Chapel Hill, NC, USA; Quantitative and Computational Biology Section, Department of Biological Sciences, University of Southern California, Los Angeles, CA, USA; Department of Molecular Physiology and Biophysics, Vanderbilt University, Nashville, TN, USA; National Health Service Blood and Transplant, Cambridge, UK; Laboratory of Statistical Immunology, Osaka University Graduate School of Medicine, Suita, Osaka, Japan

## Abstract

**SUMMARY:** Most loci identified by GWAS have been found in populations of European ancestry (EA). In trans-ethnic meta-analyses for 15 hematological traits in 746,667 participants, including 184,535 non-EA individuals, we identified 5,552 trait-variant associations at *P*<5×10^−9^, including 71 novel loci not found in EA populations. We also identified novel ancestry-specific variants not found in EA, including an *IL7* missense variant in South Asians associated with lymphocyte count *in vivo* and IL7 secretion levels *in vitro*. Fine-mapping prioritized variants annotated as functional, and generated 95% credible sets that were 30% smaller when using the trans-ethnic as opposed to the EA-only results. We explored the clinical significance and predictive value of trans-ethnic variants in multiple populations, and compared genetic architecture and the impact of natural selection on these blood phenotypes between populations. Altogether, our results for hematological traits highlight the value of a more global representation of populations in genetic studies.

## INTRODUCTION

Blood-cell counts and indices are quantitative clinical laboratory measures that reflect hematopoietic progenitor cell production, hemoglobin synthesis, maturation and release from the bone marrow, and clearance of mature or senescent blood cells from the circulation. Quantitative red blood cell (RBC), white blood cell (WBC) and platelet (PLT) traits exhibit strong heritability (*h*^2^~30-80%)(Evans et al., 1999; Hinckley et al., 2013) and have been the subject of various genome-wide association studies (GWAS), including a large study that identified >1000 genomic loci in ~150,000 individuals of European-ancestry (EA)(Astle et al., 2016).

Importantly, the distribution of hematologic traits and prevalence of inherited hematologic conditions differs by ethnicity. For example, the prevalence of anemia and microcytosis is higher among African-ancestry (AFR) individuals compared to EA individuals in part due to the presence of globin gene mutations (e.g. sickle cell, α/β-thalassemia) more common among African, Mediterranean and Asian populations (Beutler and West, 2005; Raffield et al., 2018; Rana et al., 1993). AFR individuals tend to have lower WBC and neutrophil counts partly because of the Duffy/*DARC* null variant (Rappoport et al., 2019). Among Hispanics/Latinos (HA), a common Native American functional intronic variant of *ACTN1* is associated with lower PLT count (Schick et al., 2016).

Despite these observations, non-EA populations have been severely under-represented in most blood-cell genetic studies to date (Popejoy and Fullerton, 2016; Popejoy et al., 2018; Wojcik et al., 2019). Multiethnic GWAS have been recognized as more powerful for gene mapping due to ancestry-specific differences in allele frequency, linkage disequilibrium (LD), and effect size of causal variants (Li and Keating, 2014). Since blood cells play a key role in pathogen invasion, defense and inflammatory responses, hematologic-associated genetic loci are particularly predisposed to be differentiated across ancestral populations as a result of population history and local evolutionary selective pressures (Ding et al., 2013; Lo et al., 2011; Raj et al., 2013). Given the essential role of blood cells in tissue oxygen delivery, inflammatory responses, atherosclerosis, and thrombosis (Byrnes and Wolberg, 2017; Chu et al., 2010; Colin et al., 2014; Tajuddin et al., 2016), factors that contribute to such inter-population differences in blood-cell traits may also play appreciable roles in the pathogenesis of chronic diseases and health disparities between populations.

## RESULTS

### Trans-ethnic and ancestry-specific blood-cell traits genetic associations

We analyzed genotype-phenotype associations at up to 45 million autosomal variants in 746,667 participants, including 184,424 individuals of non-EA descent, for 15 traits (**Figure 1**, **Supplementary Tables 1-4**, and **Methods**). The association results of the EA-specific meta-analyses are reported separately in a companion paper. In the trans-ethnic meta-analyses, we identified 5,552 trait-variant associations at *P*<5×10^−9^, which include 71 novel associations not reported in the EA-specific manuscript (**Supplementary Table 5**). Of the 5,552 trans-ethnic loci, 128 showed strong evidence of allelic effect heterogeneity across populations (*P*_ancestry.hetero_ <5×10−^9^) (**Supplementary Figure 1** and **Supplementary Table 5**). Ancestry-specific meta-analyses revealed 28 novel trait-variant associations (**Figure 1** and **Supplementary Tables 6-10**). However, 19 out of these 21 novel AFR-specific associations map to chromosome 1 and are associated with WBC or neutrophil counts, therefore reflecting long-range associations due to the admixture signal at the Duffy/*DARC* locus (Reich et al., 2009). We attempted to replicate all novel trans-ethnic or ancestry-specific genetic associations in the Million Veteran Program (MVP) cohort (Gaziano et al., 2016). Of the 89 variant-trait associations that we could test in MVP, 86 had a consistent direction of effect (binomial *P*=6×10^−24^), 72 had an association *P*<0.05 (binomial *P*=8×10^−79^), and 44 met the Bonferroni-adjusted significance threshold of *P*<6×10^−4^ (**Supplementary Table 11**).

**Table 1.**
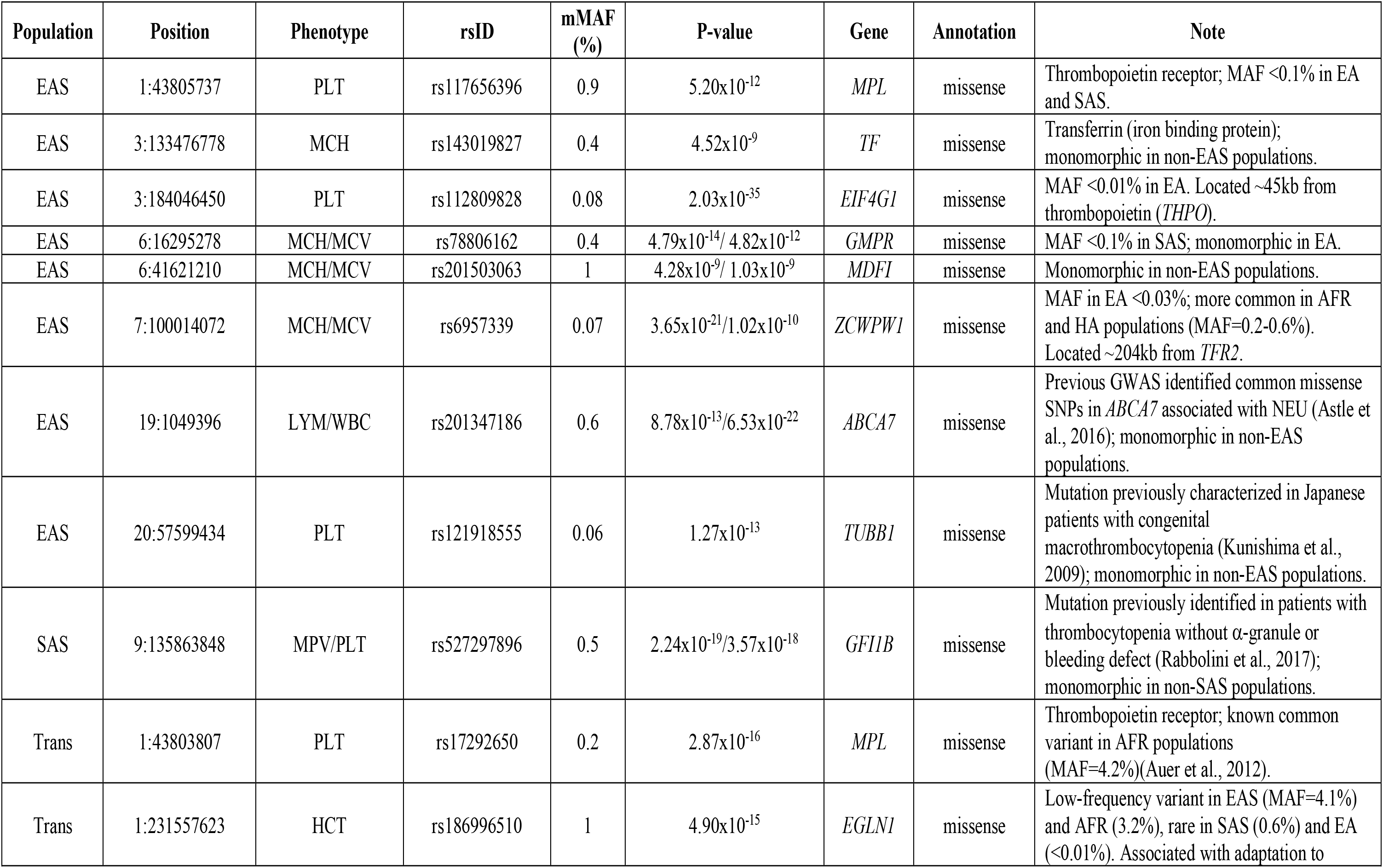
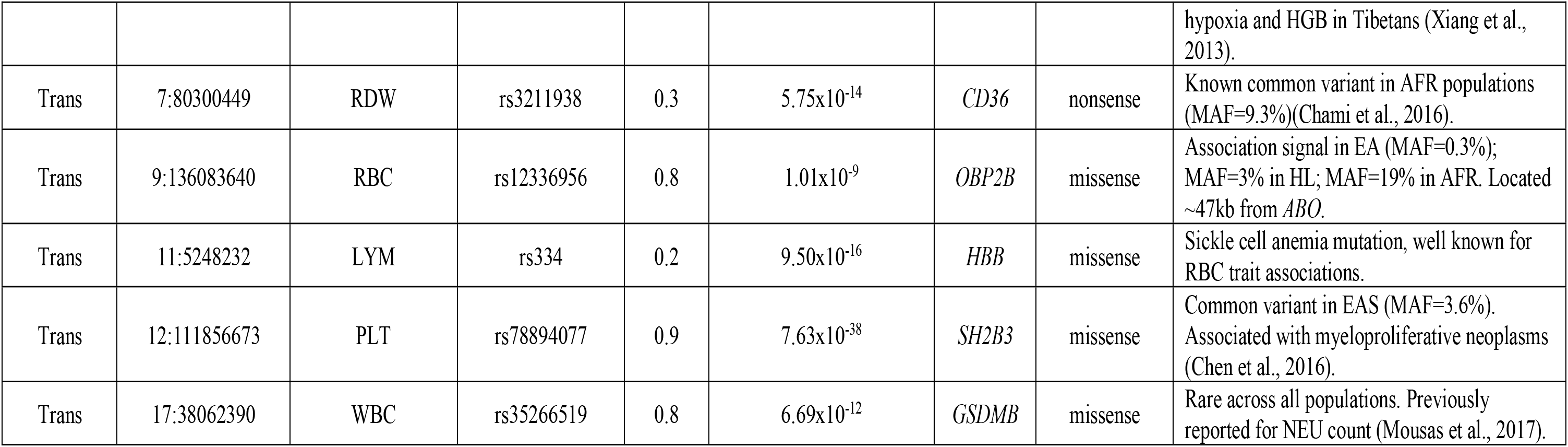
Non-synonymous variants with a minor allele frequency (MAF) ≤1% identified in non-European-ancestry (EA) populations or in the transethnic meta-analyses. The population in which each variant was discovered is listed in the first column. Complete association results for each variant are available in **Supplementary Table 15**. Genomic coordinates (chr:position) are on build hg19. For the trans-ethnic results, mMAF corresponds to the mean MAF across all studies. EA, European-ancestry; EAS, East Asians; SAS, South Asians; AFR, African-ancestry; HA, Hispanics; PLT, platelet; NEU, neutrophil; MCH, mean corpuscular hemoglobin; MCV, mean corpuscular volume; MCHC, mean corpuscular hemoglobin concentration; MPV, mean platelet volume; EOS, eosinophil; MON, monocyte; RDW, red blood cell distribution width; LYM, lymphocyte; RBC, red blood cell count; HGB, hemoglobin; WBC, white blood cell.

**Figure 1.**
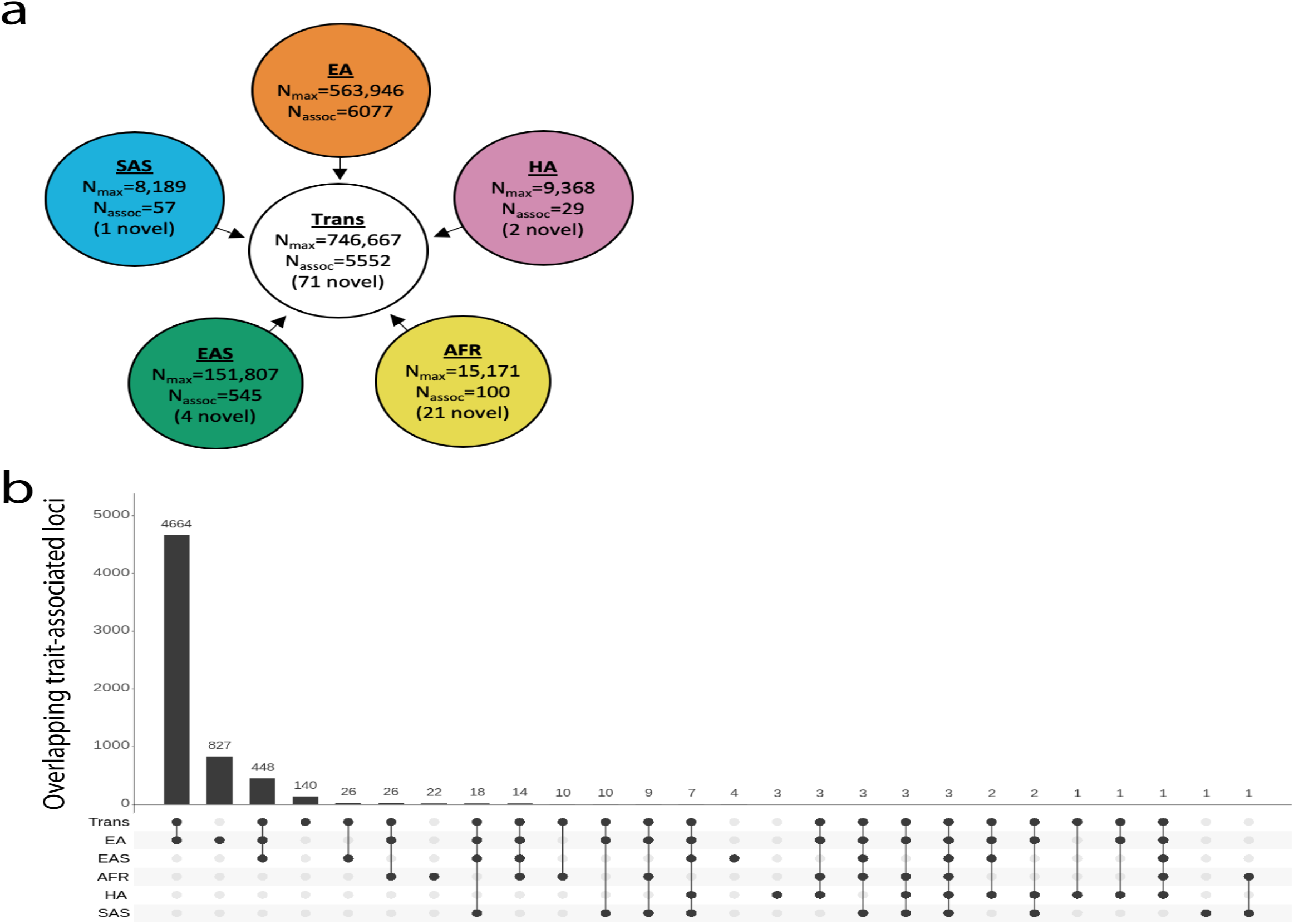
Trans-ethnic and ancestry-specific meta-analyses of blood-cell traits. (**a**) Study design of the project. We used a fixed-effect meta-analysis strategy to analyze genetic associations within each of the five populations available, and a mega-regression approach that considers allele frequency heterogeneity for the trans-ethnic association tests. (**b**) Most blood-cell trait-associated loci physically overlap between populations. Despite different sample sizes between populations, we note that few loci are found in a single population, suggesting shared genetic architecture.

For 3,552 loci with evidence of a single association signal based on conditional analyses in EA (**Supplementary Methods**), we generated fine-mapping results for each trans-ethnic or ancestry-specific dataset using an approximate Bayesian approach (**Methods**)(Wellcome Trust Case Control et al., 2012). The 95% credible sets were smaller in the trans-ethnic meta-analyses than in the EA or EAS meta-analyses (**Figure 2A**), indicating improved resolution owing to both increased sample size and different LD patterns. When comparing loci discovered in both the trans and EA analyses, we found that the 95% credible sets were 30% smaller among the trans results (median (interquartile range) number of variants per 95% credible set was 4 (2-13) in trans vs. 5 (2-16) in EA, Wilcoxon’s *P*=3×10^−4^). For instance, a locus on chromosome 9 associated with PLT counts included seven variants in the EA 95% credible set but only one in the trans set, an increase in fine-mapping resolution likely driven by limited LD at the locus in EAS (**Figure 2B**). In the trans and EA results, respectively, we identified 433 and 403 loci with a single variant in the 95% credible sets (**Figure 2C**).

**Figure 2.**
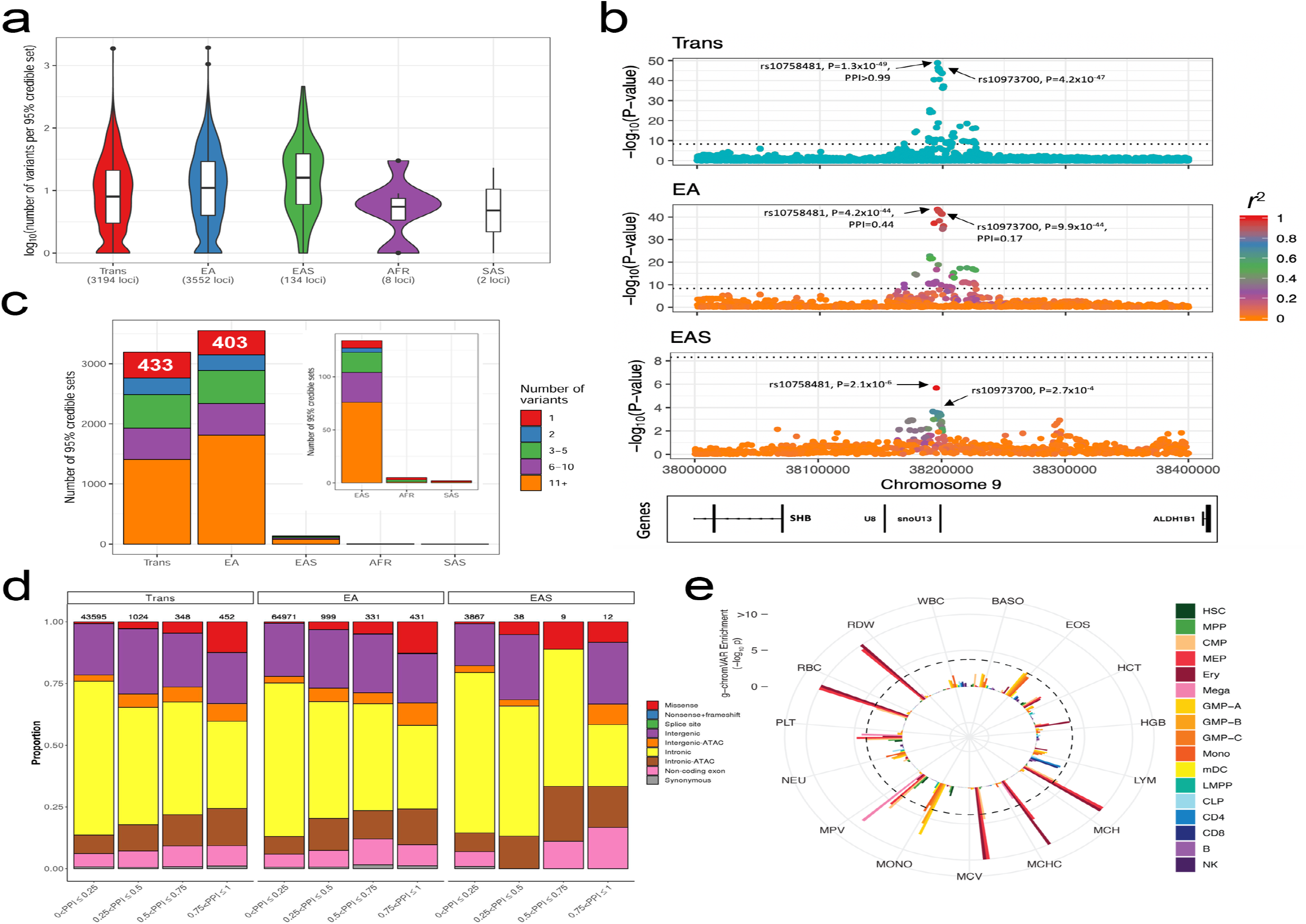
Fine-mapping of loci associated with hematological traits highlights likely functional variants. (**a**) We restricted fine-mapping to loci with evidence for a single association signal in European-ancestry (EA) populations. There are no such loci in Hispanic Americans. The 95% credible sets in the trans-ethnic meta-analyses are smaller than in the EA or East-Asian-ancestry (EAS) meta-analyses. (**b**) Trans-ethnic fine-mapping of a platelet locus. In EA individuals, the 95% credible set include 7 variants with posterior probability of inclusion (PPI) >0.04 and strong pairwise linkage disequilibrium (LD) with the sentinel variants rs10758481 (*r*^2^>0.93 in GBR from 1000 Genomes Project, middle panel). LD is similarly strong in African-, Hispanic/South American-, and South-Asian-ancestry populations from the 1000 Genomes Project. However, LD is weaker in East Asians (*r*^2^=0.68 in JPT from 1000 Genomes Project, bottom panel). In the trans-ethnic meta-analysis, rs10758481 has a PPitalic>0.99 (top panel). In EA and EAS, LD is color-coded based on pairwise *r*^2^ with rs10758481. The dotted line indicates the genome-wide significance threshold (*P*<5×10^−9^). (**c**) Number of variants in 95% credible sets in each population analyzed. In trans, EA and EAS, we identified 433, 403 and seven 95% credible sets with a single variant. (**d**) Annotation of variants in trans, EA and EAS shows a similar pattern, with a larger proportion of likely functional variants (e.g. missense, intergenic and intronic variants within ATAC-seq peaks) among variants with higher posterior probability of inclusion (PPI). (**e**) g-chromVAR results for trans variants within 95% credible sets for 15 traits. The Bonferroni-adjusted significance level (corrected for 15 traits and 18 cell types) is indicated by the dotted line. mono, monocyte; gran, granulocyte; ery, erythroid; mega, megakaryocyte; CD4, CD4+ T cell; CD8, CD8+ T cell; B, B cell; NK, natural killer cell; mDC, myeloid dendritic cell; pDC, plasmacytoid dendritic cell; MPP, multipotent progenitor; LMPP, lymphoid-primed multipotent progenitor; CMP, common myeloid progenitor; CLP, common lymphoid progenitor; GMP, granulocyte–macrophage progenitor; MEP, megakaryocyte–erythroid progenitor.

Next, we assessed our fine-mapped 95% credible sets for the presence of functional variants, which we defined as variants with coding consequences or those mapping to hematopoietic accessible chromatin. Genomic annotation of the 95% credible sets of the trans, EA and EAS hematological trait-associated loci revealed that the proportion of likely functional variants was higher among those with high PPI (**Figure 2D**). The enrichment within high-PPI categories was particularly notable for missense variants, but also observed for intronic and intergenic variants that map to open chromatin regions in precursor or mature blood cells (**Figure 2D**)(Corces et al., 2016). We used g-chromVAR to quantify the enrichment of trans, EA and EAS 95% credible set variants within regions of accessible chromatin identified by ATAC-seq in 18 hematopoietic populations (Corces et al., 2016; Ulirsch et al., 2019). We noted 22 significant trait-cell type enrichments using the trans-ethnic credible sets, all of which were lineage specific, including RBC traits in erythroid progenitors, platelet traits in megakaryocytes, and monocyte count in granulocyte-macrophage progenitors (GMP) (**Figure 2E** and **Supplementary Table 12**). Cell-type enrichments were largely consistent between fine-mapped traits found in the trans, EA and EAS loci. However, we observed two noteworthy ancestry-specific differences: the EAS results revealed significant enrichments in basophil count for the common myeloid progenitor (CMP) population and eosinophil count for the GMP population, but neither pairing reached significance in the larger EA meta-analyses (**Supplementary Figure 2**). These differences persisted even after controlling for the number of loci tested in each ancestry. This is further supported by our finding that the genetic correlations for these two traits between EA and EAS are the lowest among all studied blood phenotypes (see below).

### Phenome-wide association studies (pheWAS)

When we queried the 5,552 trans-ethnic genome-wide significant variants associated with blood-cell traits in three distinct biobanks (EA individuals from UK Biobank (UKBB), Japaneses from Biobank Japan (BBJ), African Americans from BioVU (BioVU)), we identified 1,140 phenotype-variant associations (**Supplementary Table 13**). These include 106 variants in BBJ, four variants in BioVU, and 222 variants in the UKBB. Of the four variants significant in BioVU, three were located at the β-globin locus and reflect the known clinical sequelae of sickle cell disease. Of the 1,140 associations, 246 were shared across at least two biobanks (**Methods**). Many of the associations shared between BBJ and the UKBB were related to dyslipidemia and cardiovascular diseases (**Supplementary Table 13**). Because of the large differences in sample sizes between the three biobanks (**Methods**), we reasoned that lack of power was the likely explanation for why associations were not shared across biobanks. However, it appears that differences in allele frequencies across the three primary ancestries also played a role. Overall, unique associations had greater differences in allele frequencies than shared associations (**Supplementary Figure 3**).

### Trans-ethnic predictions of hematological traits

Polygenic trait scores (PTS) developed in a single ethnically homogeneous population tend to underperform when tested in a different population (Grinde et al., 2019; Marquez-Luna et al., 2017; Martin et al., 2019). We explored whether we could combine the genome-wide significant trans-ethnic variants identified in our analyses into PTS that can predict blood-cell traits in a multi-ethnic setting. First, we used trans-ethnic effect sizes as weights to compute PTS_trans_ for each trait, and tested their performance in independent EA, AFR and HA participants from the BioMe Biobank (**Methods**). As expected because our trans-ethnic meta-analyses are dominated by EA individuals, PTS_trans_ were more predictive in EA, although their performance in HA was comparable for several traits (lymphocyte and monocyte counts, mean PLT volume)(**Supplementary Figure 4A** and **Supplementary Table 14**). Moreover, for neutrophil and WBC counts, the variance explained by the PTS_trans_ was up to three times higher in AFR and HA than in EA samples due to the inclusion of the strong Duffy/*DARC* locus (**Supplementary Figure 4A**). Because these Duffy/*DARC* variants would not have been included in PTS derived uniquely from EA association results, this illustrates an interesting feature of using trans-ethnic variants for building polygenic predictors. Next, we asked if we could increase the variance explained by calculating PTS using the same trans-ethnic variants but weighting them using ancestry-specific as opposed to trans-ethnic effect sizes. PTS_trans_ outperformed ancestry-specific PTS_AFR_ and PTS_HA_ in BioMe AFR and HA participants, respectively (**Supplementary Figure 4B-C** and **Supplementary Table 14**). This result likely indicates that the discovery sample size for these two populations is still too small to provide robust estimates of the true population-specific effect sizes.

### Rare coding blood-cell-traits-associated variants

The identification of rare coding variants has successfully pinpointed candidate genes for many complex traits, including blood-cell phenotypes (Auer et al., 2014; Chami et al., 2016; Eicher et al., 2016; Justice et al., 2019; Marouli et al., 2017; Mousas et al., 2017; Tajuddin et al., 2016). Our trans-ethnic and non-EA ancestry-specific meta-analyses yielded 16 coding variants with minor allele frequency (MAF) <1% (**Table 1** and **Supplementary Table 15**). This list includes variants of clinical significance (variants in *TUBB1*, *GFI1B*, *HBB*, *MPL* and *SH2B3*) and variants that nominate candidate genes within GWAS loci (*ABCA7, GMPR*) (**Table 1**). Our analyses also retrieved a known missense variant in *EGLN1* (rs186996510) that is associated with high-altitude adaptation and hemoglobin levels in Tibetans (Lorenzo et al., 2014; Xiang et al., 2013). We noted a missense variant in *IL7* (rs201412253, Val18Ile) associated with increased lymphocyte count in South Asians (SAS)(*P*=4.4×10^−10^) (**Figure 3A** and **Supplementary Table 16**). This variant is low-frequency in SAS (MAF=2.6%) but rare in other populations (MAF <0.4%). IL7 encodes interleukin-7, a cytokine essential for B- and T-cell lymphopoiesis (Lin et al., 2017). IL7 is synthesized as a proprotein that is cleaved prior to secretion, and the *IL7*-Val18Ile variant localizes to the IL7 signal peptide comprising the first 25 amino acids. To determine if this variant alters IL7 secretion, we engineered HEK293 cells with either IL7 allele (**Methods**). Although there was no difference in *IL7* RNA expression levels (*t*-test *P*=0.63), we found that the IL7-18Ile allele, which associates with higher lymphocyte counts in SAS individuals, significantly increased IL7 protein secretion in this heterologous cellular system (+83%, *P*=2.7×10^−5^)(**Figure 3B**). Unfortunately, the relatively small sample size of the SAS cohort prevented us from testing whether this *IL7* variant may be associated with other relevant phenotypes, such as cancer risk or susceptibility to infections (Lin et al., 2017).

**Figure 3.**
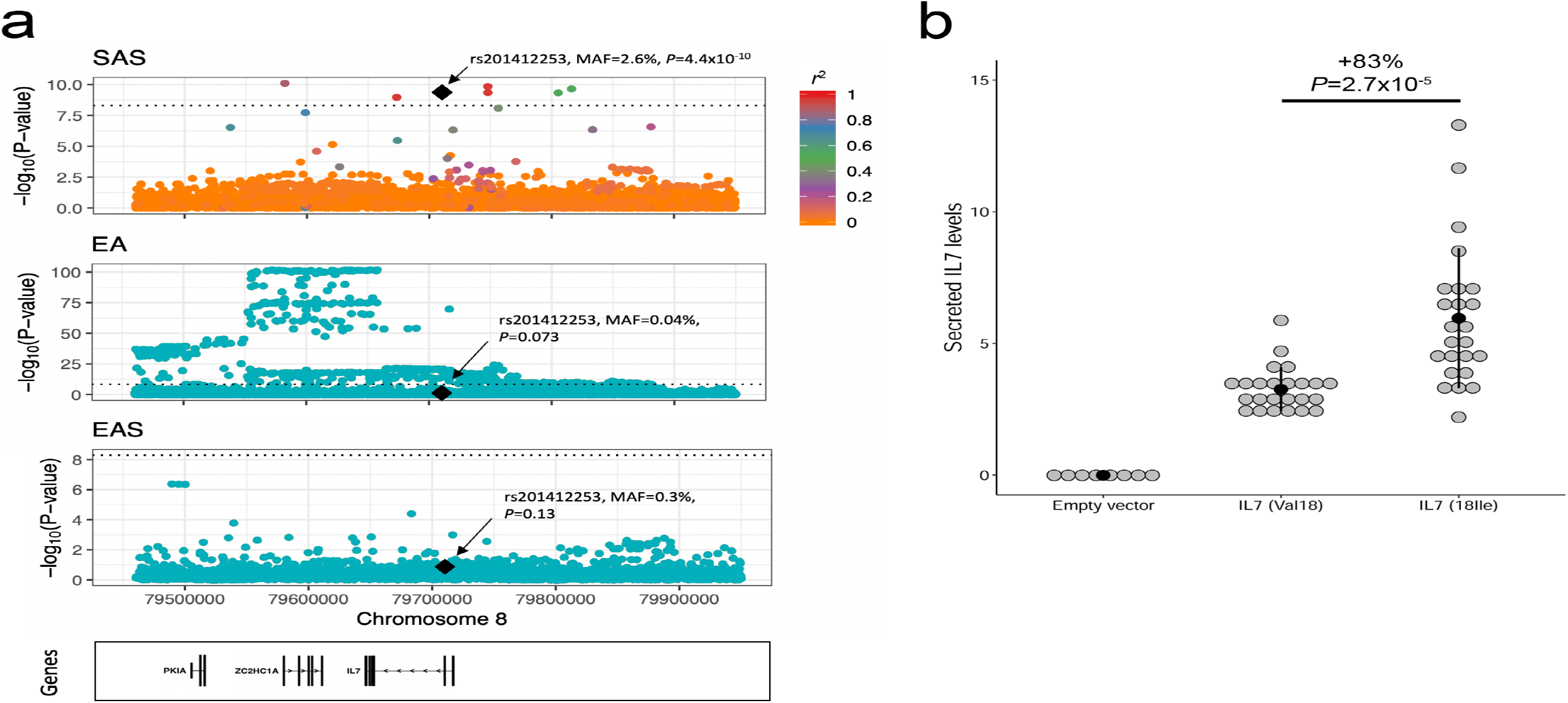
A South-Asian-ancestry *IL7* missense variant associates with increased lymphocyte count in humans and IL7 secretion in an heterologous cellular system. (**a**) Lymphocyte count association results at the *IL7* locus in South Asians (SAS), European-ancestry participants (EA) and East Asians (EAS). In SAS, there are 7 genome-wide significant variants near *IL7*, but only rs201412253 is coding. Linkage disequilibrium (LD) *r*^2^ is from 1000 Genomes Project SAS populations. In EA, the sentinel variant is located downstream of *IL7*; rs201412253 is rare (minor allele frequency=4×10^−4^) and not significant (*P*=0.073). In EAS, the locus is not associated with lymphocyte count. rs201412253 is monomorphic in 1000 Genomes Project EA and EAS so we could not calculate pairwise LD. (**b**) The 18Ile allele at *IL7*-rs201412253 increases IL7 secretion in a heterologous cellular system. Our ELISA assay did not detect secreted IL7 in clones generated with an empty vector. We tested eight independent clones for each *IL7* alleles. Each experiment was done in duplicate, and we performed the experiments three times (grey circles). The black dots and vertical lines indicate means and standard deviations. We assess statistical significance by linear regression correcting for experimental batch effects.

**Figure 4.**
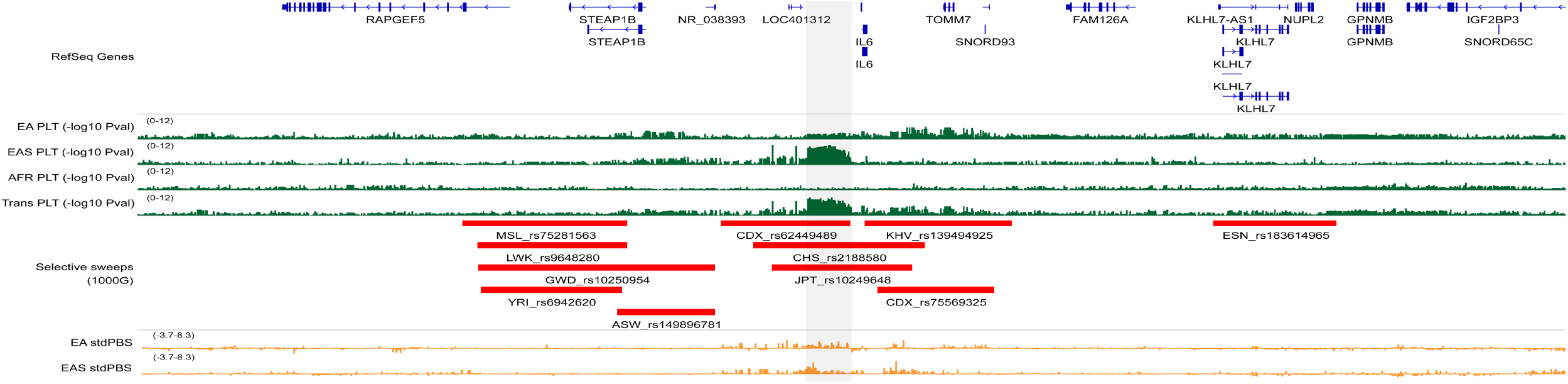
Selective sweep and association with platelet count at the *IL6* locus in East Asians. The grey rectangle highlights a genomic region upstream of *IL6* that is strongly associated with platelet (PLT) count. This association signal is driven by results from East Asians (EAS), and is absent from other populations, including European-(EA) and African-ancestry (AFR) individuals (green). The region overlaps several selective sweeps detected in EAS from the 1000 Genomes Project (CDX, CHS, JPT, KHV). In orange, we provide standardized population branch site (stdPBS) metrics in EA and EAS, indicative of allele frequency differentiation at this locus between these two populations.

### Genetic architecture of blood-cell traits in EA and EAS populations

We used several different approaches to quantify similarities and differences in genetic architecture of hematologic traits across populations. Focusing on the two largest studied populations, EA and EAS, we calculated heritability for all blood traits and found them to be highly concordant between ancestries (Pearson’s *r*=0.75, *P*=0.0033)(**Supplementary Table 17**)(Bulik-Sullivan et al., 2015b). Likewise, within-ancestry genetic correlation coefficients (*r*_g_) between pairs of hematological traits were highly concordant across ancestries (Pearson’s *r*=0.97, *P*<2.2×10^−16^)(**Supplementary Figure 5**)(Bulik-Sullivan et al., 2015a). We then used the Popcorn method to directly measure genetic correlations for blood-cell traits between EA and EAS using summary statistics for common variants (Brown et al., 2016). For all 13 traits available in both EA and EAS, genetic correlations were high (lowest for basophils (*r*_g_=0.30) and highest for MCH (*r*_g_=0.66)), but significantly different than 1 (*P*<3×10-6)(**Supplementary Table 18**). This suggests that even when considering only common variants, the genetic architecture of blood phenotypes is substantially different between populations.

### Natural selection at blood-cell trait loci

Natural selection can account for differences in association results between populations, as highlighted by our analyses of rare coding variants which includes several loci known to be under selection (*CD36*, *HBB*, *EGLN1*)(**Table 1**). To further explore this possibility, we assessed whether variants that tag selective sweeps (tagSweeps, variants with the highest integrated haplotype score (iHS)) within continental populations from the 1000 Genomes Project (1000G) are associated with blood-cell phenotypes (Johnson and Voight, 2018). We found a genome-wide enrichment of associations results between tagSweeps and hematological traits, particularly within EA, EAS and AFR populations (**Supplementary Figure 6** and **Supplementary Table 19**). To rule out simple overlaps due to the large number of sweeps and blood-cell trait loci, we compared the number of genome-wide significant tagSweeps in EA, EAS and AFR with the number of significant variants among 100 sets of matched variants (**Methods**). We found significant enrichment of selective sweeps for WBC (EA, EAS, AFR), monocytes (EA, AFR), eosinophils (EA), neutrophils (AFR), lymphocytes (EAS), and PLT (EA, EAS)(**Supplementary Table 20**).

In AFR and HA, the enrichments for WBC, neutrophils and monocytes were entirely driven by selective sweeps on chromosome 1 near Duffy/*DARC* (Reich et al., 2009). Only three additional loci shared evidence of associations with blood-cell traits and positive selection across populations: HLA, *SH2B3* (Zhernakova et al., 2010) and *CYP3A5* (Chen et al., 2009). We found eight and 100 non-overlapping selective sweeps with variants associated with hematological traits in EAS and EA, respectively (**Supplementary Table 21**). Six of the eight EAS-specific tagSweeps are also associated with blood-cell traits in EA participants, indicating that these regions do not account for population differences in hematological trait regulation (**Supplementary Table 21**). One of the remaining two variants is located at the *HBS1L-MYB* locus and, although it is not associated with blood-cell traits in EA, there are many other variants near *MYB* associated with blood phenotypes in EA (**Supplementary Table 6**). The remaining selective sweep highlighted by this analysis is located upstream of *IL6* (**Figure 4**). The tagSweep at this locus, rs2188580, is strongly associated with PLT count in EAS (*P*_EAS_=2.8×10-9, *P*_EA_=0.0022), is differentiated between EAS and EA as indicated by the population branch statistic (PBS)(Yi et al., 2010)(C-allele frequency in EAS=44%, 4% in EA; standardized PBS_EAS_=7.353), and overlaps selective sweeps identified in several EAS populations from the 1000G (e.g. iHS_CHS_=3.935)(**Figure 4**). The *IL6* locus has previously been associated with WBC traits in EA (Astle et al., 2016), but our finding is the first report of its association with PLT. *IL6* encodes interleukin-6, a cytokine that is a maturation factor for megakaryocytes, the precursors of PLT (Kimura et al., 1990). Further supporting the role of IL6 signaling in PLT biology, a well-characterized missense variant in the IL6 receptor gene (*IL6R*-rs2228145)(van Dongen et al., 2014) is also associated with PLT count in EAS (*P*=4.3×10^−6^).

## DISCUSSION

Our meta-analyses of 15 hematological traits in up to 746,667 individuals represents one of the largest genetic study of clinically relevant complex human traits across diverse ancestral groups. We have continued to expand the repertoire of loci and genes that contribute to interindividual variation in blood-cell traits, with potential implications for hematological diseases, but also other conditions such as cancer, immune and cardiovascular diseases. Our results hold a number of implications for future human genetic studies. First, we showed that adding even a “modest” number of non-EA participants to GWAS can yield important biology, such as the identification of a lymphocyte count-associated *IL7* missense variants in 8,189 South Asians (**Figure 3**). Second, loci that underlie variation in blood-cell traits represent a broad mixture of shared associations (i.e. similar allele frequencies and effect sizes across populations) and heterogeneous associations (i.e. dissimilar allele frequencies and effect sizes across populations). This result contributes to mounting evidence that a full accounting of the genetic basis of complex human traits will require a thorough catalog of global genetic and phenotypic variation. Third, because of heterogeneity across populations in both allele frequencies and patterns of LD, fine-mapping of association signals can be substantially aided by including multiple ancestries. This will have a dramatic impact on the success of large-scale efforts aimed at functionally characterizing GWAS findings. As more studies seek to unravel the causal variants that underlie complex traits associations, we anticipate that genetic evidence from diverse ancestries will play an important role.

## SUPPLEMENTERY INFORMATION

**Supplementary Information** is linked to the online version of the paper.

## ACKNOWLEDGMENTS

We thank all participants, as well as Dr. John D. Rioux for providing the *IL7* ORF. A full list of acknowledgments appears in the **Supplementary Information**. Part of this work was conducted using the UK Biobank resource (Projects number 11707 and 13745).

## AUTHOR CONTRIBUTIONS

### Writing Group (wrote and edited manuscript)

Ming-Huei Chen, Laura M. Raffield, Abdou Mousas, Erik L. Bao, Alexander P. Reiner, Paul L. Auer, Guillaume Lettre. All authors contributed and discussed the results, and commented on the manuscript.

### Data preparation group (checked and prepared data from contributing cohorts for meta-analyses and replication)

Ming-Huei Chen, Laura M. Raffield, Abdou Mousas, Jennifer E. Huffman, Guillaume Lettre, Paul L. Auer.

### Meta-analyses (discovery and replication)

Ming-Huei Chen, Laura M. Raffield, Abdou Mousas, Saori Sakaue, Jennifer E. Huffman, Tao Jiang, Parsa Akbari, Dragana Vuckovic, Erik L. Bao, Arden Moscati, Ken Sin Lo, Caleb A. Lareau, Michael H. Guo, Tim Kacprowski, Fotis Koskeridis, Ani Manichaikul, Michael Preuss, Cassandra N. Spracklen.

### Fine-mapping and functional annotation

Abdou Mousas, Ming-Huei Chen, Laura M. Raffield, Erik L. Bao, Caleb A. Lareau, Ken Sin Lo, Vijay G. Sankaran, Paul L. Auer, Guillaume Lettre.

### pheWAS, polygenic prediction, genetic architecture and natural selection

Saori Sakaue, Arden Moscati, Regina Manansala, Minhui Chen, Ken Sin Lo, Huijun Qian, Yun Li, Charleston W.K. Chiang, Ruth J.F. Loos, Alexander P. Reiner, Guillaume Lettre, Paul L. Auer.

### Functional characterization of IL7

Mélissa Beaudoin, Véronique Laplante, Guillaume Lettre, Jean-François Gauchat

## AUTHOR INFORMATION

Summary genetic association, fine-mapping and g-chromVAR results are available online: http://www.mhi-humangenetics.org/en/resources. Competing financial interests are declared in the

**Supplementary Information**. Correspondence and requests should be addressed to P.L.A or G.L.

## METHODS

### Study design and participants

All participants provided written informed consent and the project was approved by each institution’s ethical committee. **Supplementary Table 1** lists all participating cohorts. The SNPs we identified are available from the NCBI dbSNP database of short genetic variations (https://www.ncbi.nlm.nih.gov/projects/SNP/). No statistical methods were used to predetermine sample size. The experiments were not randomized and the investigators were not blinded to allocation during experiments and outcome assessment.

### Phenotypes

Complete blood count (CBC) and related blood indices were analyzed as quantitative traits. The descriptive statistics for each phenotype in each cohort analyzed are in **Supplementary Table 2**. Exclusion criteria and phenotype modeling in the UK Biobank (UKBB)(European-ancestry individuals), INTERVAL, and Biobank Japan (BBJ) have been described previously (Astle et al., 2016; Kanai et al., 2018). For all other studies, we followed the protocol developed by the Blood-Cell Consortium (Chami et al., 2016; Eicher et al., 2016; Tajuddin et al., 2016). Briefly, we excluded when possible participants with blood cancer, acute medical/surgical illness, myelodysplastic syndrome, bone marrow transplant, congenital/hereditary anemia, HIV, end-stage kidney disease, splenectomy, and cirrhosis, as well as pregnant women and those undergoing chemotherapy or erythropoietin treatment. We also excluded extreme blood-cell measures: WBC>200×10^9^ cells/L, HGB>20 g/dL, HCT>60%, and PLT>1000×10^9^ cells/L. For WBC subtypes, we analyzed log_10_-transformed absolute counts obtained by multiplying relative counts with total WBC count. For all phenotypes in all studies, we corrected the blood-cell phenotypes for sex, age, age-squared, the 10 first genetic principal components, and other cohort-specific covariates (e.g. recruitment center) using linear regression analysis. We applied rank-based inverse normal transformation to the residuals form the regression analysis and used the normalized residuals to test for association with genetic variants.

### Genotype quality-control and imputation

The genotyping array and quality-control steps used by each cohort as well as their quality-control steps are listed in **Supplementary Table 3**. Unless otherwise specified, all studies applied the following criteria: samples were removed if the genotyping call rate was <95%, if they showed excess heterozygosity, if we identified gender mismatches or sample duplicates, or if they appeared as population outliers in principal component analyses nested with continental populations from the 1000 Genomes Project (Genomes Project et al., 2012). We removed monomorphic variants, as well as variants with Hardy-Weinberg *P*<1×10^−6^ and call rate <98%.

Genotype imputation for the UKBB, INTERVAL, and BBJ have been described in details elsewhere (Astle et al., 2016; Bycroft et al., 2018; Kanai et al., 2018). For all other studies, unless specified in **Supplementary Table 3**, we applied the following steps for genotype imputation of autosomal variants. We aligned all alleles on the forward strand of build 37/hg19 of the human reference genome (http://www.well.ox.ac.uk/~wrayner/strand) and converted files into the VCF format. We then applied checkVCF (http://genome.sph.umich.edu/wiki/CheckVCF.py) to confirm strand and allele orientation. We carried out genotype imputation using the University of Michigan (https://imputationserver.sph.umich.edu) or the Sanger Institute (https://imputation.sanger.ac.uk/) imputation servers. We phased genotype data using SHAPEIT (Delaneau et al., 2013), EAGLE (Loh et al., 2016), or HAPI-UR (Williams et al., 2012). For populations of European ancestry, we used reference haplotypes from the Haplotype Reference Consortium (HRC r1.1 2016) for imputation (McCarthy et al., 2016), whereas reference haplotypes from the 1000 Genomes Project (Phase 3, Version 5)(Genomes Project et al., 2012) were used for non-European ancestry participants.

### Study-level statistical analyses

We tested an additive genetic model of association between genotype imputation doses and inverse normal transformed blood-cell phenotypes. We analyzed the major ancestry groups (European (EA), East Asian (EAS), African (AFR), Hispanic-Latino (HL), South Asian (SAS)) separately and used linear mixed-effect models implemented in BOLT-LMM (Loh et al., 2018), EPACTS (https://genome.sph.umich.edu/wiki/EPACTS), or EMMAX (Kang et al., 2010) to account for cryptic and known relatedness. Autosomal single nucleotide variants were analyzed in all contributing studies. For simplicity, we only analyzed insertion-deletion (indel) variants from UKBB and INTERVAL, since a similar reference panel was used for genotype imputation.

### Centralized quality-control and meta-analyses

We performed a centralized quality-control check on the association results of each single study using EasyQC (v9.0)(Winkler et al., 2014). By mapping variants of each study to the appropriate ethnicity reference panel (HRC for EA and 1000 Genomes Project Phase3 for non-EA participants), we were able to harmonize alleles and markers across all studies. We were also able to assess the presence of flipped alleles per study and check for excessive allele frequency discrepancies using allele frequency reference data. We also inspected quantile-quantile (QQ) plots generated by EasyQC and the corresponding genomic inflation factors as well as SE-N plots (inverse of the median standard error vs. the square root of the sample size) to evaluate potential issues with, for example, trait transformation or unaccounted relatedness. We removed variants with imputation quality metric (INFO score) ≤0.4. Except for three studies, we also removed variants with minor allele count (MAC) ≤5. For UKBB EA, Women Health Initiative (WHI), and GERA (EA), we instead applied a MAC ≤20 filter because empirical observations suggested that unusual inflation of the test statistics (i.e. extreme effect sizes and standard errors) was due to rarer variants. To simplify handling of tri-allelic and indel variants, which have the same genomic coordinates but different alleles, we created a unique variant ID for each tested variant. Specifically, we assigned a chromosome:position(hg19)_allele1_allele2 unique ID to each variant, in which the order of the allele in the ID was based on the lexicographical order or the indel length. We performed inverse variance-weighted fixed-effect meta-analyses with GWAMA (v2.2.2)(Magi and Morris, 2010) and trans-ethnic meta-analyses with MR-MEGA (v0.1.5)(Magi et al., 2017). For MR-MEGA, we calculated four axes of genetic variation, the default recommendation, to separate global population groups.

### Statistical significance, genomic inflation and locus definition

For each meta-analysis, we calculated the genomic inflation factor (λ_GC_) for all variants, which were modest when considering the large sample sizes (λ_GC_ range: 0.9-1.2) (**Supplementary Table 4**). We used α ≤5×10^−9^ after GC-correction to declare statistical significance, accounting for the inflation of the test statistics and the number of blood-cell traits analyzed. To count the number of loci that we discovered, we first identified the most significant variants (with P≤5×10^−9^) and extended the physical region around that variant 250-kb on each side. Overlapping loci were merged, and we used the most significant variant within the interval as the sentinel variant. In this manuscript, we defined as novel a locus if no variants were previously reported in the literature to be associated with the specific blood-cell trait and if the locus is not reported in the companion manuscript that focuses on EA-specific genetic discoveries.

### Million Veteran Program (MVP) blood-cell trait analyses for replication

#### Phenotyping

Phenotyping methods published by the EMERGE Consortium and available on PheKB (https://phekb.org/) were used for retrieving lab data and exclusion criteria for all blood cell indices. This information was pulled from the VA electronic medical records for all MVP participants. Lab data was subject to the Boston Lab Adjudication Protocol. This entails five steps: (i) compile an initial spreadsheet of possible relevant lab tests, (ii) Subject Matter Expert (SME) does an initial review of possible tests, (iii) analyst adds relevant LOINC codes for SME review, (iv) second Subject Matter Expert (SME) review, (v) creation of a Lab Phenotype Table/Data Set. After restricting to only outpatient labs and applying the EMERGE exclusion criteria, for each trait and each person, the minimum, maximum, mean, median, SD, and number of labs was recorded. Values were compared to those from UKBB (Astle et al., 2016).

#### Genotyping

DNA extracted from whole blood was genotyped using a customized Affymetrix Axiom biobank array, the MVP 1.0 Genotyping Array. With 723,305 total DNA sequence variants, the array is enriched for both common and rare variants of clinical importance in different ethnic backgrounds (Klarin et al., 2018).

#### Analysis

The median lab value was the trait used for analysis. Linear regression models were run under an additive model in plink2 on 1000G (v3p5) imputed dosages. Analyses were run using models described above within each race/ethnicity stratum (AFR, ASN, EA, HA) classified based on their genotype data using HARE (Fang et al., 2019). Meta-analyses for the trans-ethnic analyses were completed in METAL (Willer et al., 2010).

### Heritabilities and genetic correlations

We calculated heritabilities and genetic correlations between blood-cell traits within the EA and EAS populations using default parameters implemented in the LD score regression method (**Supplementary Table 17** and **Supplementary Figure 5**)(Bulik-Sullivan et al., 2015a; Bulik-Sullivan et al., 2015b). For genetic correlation of the same phenotype between ancestral populations, we used Popcorn (Brown et al., 2016). Briefly, Popcorn uses a Bayesian framework to estimate, using genome-wide summary statistics, the genetic correlation of the same phenotype but in two different populations (in our case, between EA and EAS). It reports the trans-ethnic genetic-effect correlation (ρ_ge_), i.e. the correlation coefficient of per-allele SNP effect sizes, but also the trans-ethnic genetic impact correlation (ρ_gi_), which includes a normalization of the effect based on allele frequency (**Supplementary Table 18**). To address whether a difference in the sample size for the EA and EAS meta-analyses could impact the Popcorn results, we repeated our analyses using the current EAS results (N_max_=151,807) and EA results from preliminary analyses of the UKBB dataset (N_max_=87,265)(Astle et al., 2016). These analyses confirmed that for common variants, cross-ancestry EA-EAS genetic correlations are significantly different (but non-null). Both LD score regression and Popcorn are not amenable to admixed populations, and cannot handle rare variants. For these reasons, we limited these analyses to the large EA and EAS populations and focused on common variants from the 1000 Genomes Project.

### Statistical fine-mapping

No fine-mapping methods currently exist to handle admixed populations. Furthermore, for some of the ethnic groups analyzed here, we did not have access to a sufficiently large reference panel to properly account for LD, complicating conditional analyses and fine-mapping efforts. For these reasons, we fine-mapped the ancestry-specific fixed-effect meta-analyses by adapting the method proposed by Maller et al. (Wellcome Trust Case Control et al., 2012) in order to assign posterior probability of inclusion (PPI) to each variant and construct 95% credible sets.

This method makes the strong assumption that there is a single independent causal variant at the tested locus. For this reason, we limited our Bayesian fine-mapping to loci where we identified a single independent association signal by conditional analysis in EA individuals from the UKBB (**Supplementary Methods**). Because EA represented the largest group, we then inferred that there was also a single association signal in the other populations at these loci, an inference that may not always be right. Briefly, we added 250-kb on either side of genome-wide significant variants (P<5×10^−9^) and merged loci when they overlapped. For the loci identified in the ancestry-specific meta-analyses, we converted P-values into approximate Bayes factors (aBF) using (Wakefield, 2009; Wellcome Trust Case Control et al., 2012):

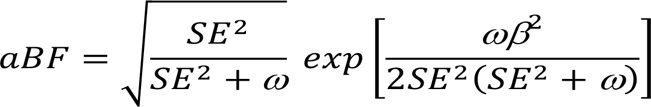

where β and SE are the variant’s effect size and standard error, respectively, and ω denotes the prior variance in allelic effects, taken here to be 0.04 (Wakefield, 2007). For the trans-ethnic results, we directly used Bayes factors calculated by MR-MEGA (Magi et al., 2017). We calculated PPI of each variant by dividing the variant’s aBF by the sum of the aBF for all the variants within the locus. We generated the 95% credible sets by ordering all variants in a given locus from the largest to the smallest PPI and by including variants until the cumulative sum of the PPI ≥95% (Mahajan et al., 2018). All variants that map to 95% credible sets are available online (see **URL**).

### Functional annotation

To derive basic functional annotation information, we annotated all variants included in 95% credible sets from ancestry-specific and trans-ethnic meta-analyses with the Variant Effect Predictor (VEP)(https://useast.ensembl.org/info/docs/tools/vep/index.html), compiling both all consequences and the most severe consequence for Ensembl/GENCODE transcripts. We also specifically annotated rare coding variants using VEP (defined as any variant with MAF <1% in a given analysis, with a GC-corrected P-value <5×10^−9^, and annotated as a missense_variant, stop_gained, stop_lost, splice_donor, or a splice_acceptor, regardless of fine-mapping results). We removed all variants with a GC-corrected P-value <5×10^−9^ in EA, in the MHC region, and, in analyses including individuals with at least some African ancestry, on chromosome 1 for neutrophils and total WBC count and for RBC traits near the chromosome 11 β-globin and the chromosome 16 α-globin loci.

Bias-corrected enrichment of blood trait variants for chromatin accessibility of 18 hematopoietic populations was performed using g-chromVAR, which has been previously described in detail (Ulirsch et al., 2019). In brief, this method weights chromatin features by fine-mapped variant posterior probabilities and computes the enrichment for each cell type versus an empirical background matched for GC content and feature intensity. For chromatin feature input, we used a consensus peak set for all hematopoietic cell types with a uniform width of 500 bp centered at the summit. For variant input, we included all fine-mapped variants within 95% credible sets of the trans-ethnic GWAS. We also ran g-chromVAR for each ancestry-specific meta-analysis, keeping all other parameters the same, but using fine-mapped variants with the 95% credible sets of each ancestry-specific study. Finally, to control for the number of loci tested within each ancestry-specific study, we first ranked the loci of the largest cohort (i.e. EA) by sentinel variant p-value, and then subset only the top *n* loci, where *n* equals the number of loci in the smaller cohort (e.g. EAS) for the same trait. We then ran g-chromVAR on the subset of variants falling within these top *n* loci.

### Phenome-wide association study (pheWAS) analysis

#### UK Biobank (UKBB)

We extracted pheWAS results for a list of 5552 variants in UKBB ICD PheWeb hosted at the University of Michigan (Accessed 21 August 2019). To account for severe imbalance in case-control ratios, we selected the output from the SAIGE analyses (http://pheweb.sph.umich.edu/SAIGE-UKB/) based on 408,961 samples from White British participants (Zhou et al., 2018). In total, 1403 phecodes were tested for association. All results were downloaded using R, and were parsed and organized into data table format using the data.table, rvest, stringr, dplyr and tidyr packages.

#### Biobank Japan (BBJ)

We performed a pheWAS for the lead variants identified by the trans-ethnic meta-analyses. From the list of all the significantly associated variants with blood cell-related traits, we extracted those genotyped or imputed in the BBJ project (*n*_SNP_ = 4,255). Next, we curated the phenotype record of the disease status and clinical values for the same individuals analyzed in the discovery phase (*n*_indiv_ = 143,988). Then, we performed the logistic regression analyses for 22 binary traits (20 diseases and 2 behavioral habits) which had a sufficient number of case samples (*n*_case_ = 2,500). Regression models were adjusted for age, sex and 20 principal components as covariates. Trait-specific covariates are described elsewhere (Kanai et al., 2018).

#### BioVU

BioVU is the biobank of Vanderbilt University Medical Center (VUMC) that houses de-identified DNA samples linked to phenotypic data derived from electronic health records (EHRs) system of VUMC. The clinical information is updated every 1-3 months for the de-identified EHRs. Detailed description of program operations, ethical considerations, and continuing oversight and patient engagement have been published (Roden et al., 2008). DNA samples were genotyped with genome-wide arrays including the Multi-Ethnic Global (MEGA) array, and the genotype data were imputed into the HRC reference panel(McCarthy et al., 2016) using the Michigan imputation server (Das et al., 2016). Imputed data and the 1000 Genome Project data were combined to carry out principal component analysis (PCA) and African-American samples were extracted for analysis based on the PCA plot. PheWAS were carried out for each SNP with the specified allele (Denny et al., 2010). Phenotypes were derived from billing codes of EHRs as described previously (Carroll et al., 2014). Each phenotype (‘phecode’) has defined case, control and exclusion criteria. We required two codes on different visit days to instantiate a case for each phecode. In total, 1815 phecodes were tested for association. Association between each binary phecode and a SNP was assessed using logistic regression, while adjusting for covariates of age, sex, genotyping array type/batch and 10 principal components of ancestry.

#### Merging across biobanks

We defined statistical significance within each biobank to be a Bonferroni corrected level of 0.05/*pq*, where *p* is the number of phecodes tested and *q* is the number of variants tested. We merged results across UKBB and BioVU by matching on phecode, as these two biobanks used the same phecode system for classifying outcomes. To merge with BBJ, we cross-referenced the 22 outcomes in BBJ with the phecode library used by BioVU/UKBB. Matches were determined based on phenotype similarity between the BioVU/UKBB phenotype description and the outcomes described in Nagai et *al.* (Nagai et al., 2017).

### Polygenic trait score (PTS) analyses

We restricted these analyses to variant-trait associations that reached genome-wide significance (*P*<5×10^−9^) in the trans-ethnic MR-MEGA meta-analyses (**Supplementary Table 5**). For each of these variant-trait pairs, we calculated an effect size – hereafter referred to as trans weights – using the fixed-effect meta-analysis method implemented in GWAMA and all cohorts available (Magi and Morris, 2010). For the same variants, we also retrieved the ancestry-specific effect sizes (or weights). We calculated the PTS using plink2 by summing up the number of trait-increasing alleles (or imputation doses) that were weighted by their corresponding trans (PTS_trans_) or ancestry-specific (PTS_EA_, PTS_AFR_, PTS_HA_) weights. The variance explained by the PTS on corrected and normalized blood-cell traits was calculated in R using linear regression. For these analyses, we had access to 2,651 AFR, 5,048 EA and 4,281 HA BioMe participants, as well as 2546 AFR ARIC participants, that were not used in the discovery effort.

### Analysis of natural selection

To quantify the contribution of positive selection on blood-cell trait variation, we used the recent map of selective sweeps identified in the different populations of the 1000 Genomes Project (Johnson and Voight, 2018). We grouped the sweeps identified in the 26 1000 Genomes Project populations into five larger populations that correspond to our ancestry-specific meta-analyses: Europe-ancestry (CEU, TSI, GBR, FIN, IBS); East-Asian-ancestry (CHB, JPT, CHS, CDX, KHV); African-ancestry (YRI, LWK, GWD, MSL, ESN, ASW, ACB); South-Asian-ancestry (GIH, PJL, BEB, STU, ITU); and Hispanic/Latino-ancestry (MXL, PUR, CLM, PEL). Following the nomenclature by Johnson and Voight(Johnson and Voight, 2018), each selective sweep is summarized by the variant located within the sweep that has the highest iHS value. iHS (Integrated Haplotype Score) is a statistic to quantify evidence of recent positive selection. A high positive iHS score (iHS > 2) means that haplotypes on the ancestral allele background are longer compared to derived allele background. A high negative iHS score (iHS < −2) means that the haplotypes on the derived allele background are longer compared to the haplotypes associated with the ancestral allele. We retrieved the blood-cell trait association results for these sweep-tagging SNPs from the ancestry-specific meta-analyses (**Supplementary Table 19**). To determine if the inflation observed in the QQ plots was significant, we generated 100 sets of SNPs that match the selective sweep-tagging SNPs based on allele frequency, gene proximity, and the number of LD proxies in European-ancestry, East-Asian-ancestry and African-ancestry individuals using SNPsnap (Pers et al., 2015). For these analyses, we excluded the HLA region and variants in LD (*r*^2^>0.5). We computed empirical significance by tallying the number of sets with the same or more genome-wide significant variants than the canonical sets of selective sweep-tagging SNPs (**Supplementary Table 20**).

We also computed the population branch statistic (PBS)(Yi et al., 2010). PBS measures the amount of allele frequency change in the population since its divergence from the other two populations. For a target population, PBS is calculated as:

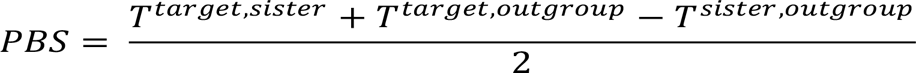

where *T* = − log(1 − *F*_ST_) is an estimate of the divergence time between two populations. Here, F_ST_ between each pair of populations was estimated using Weir and Cockerham’s estimate (Weir and Cockerham, 1984). We then divided all variants with calculated PBS into 50 bins of equal size by derived allele count in the target population, and then standardized the raw PBS values within each bin. To calculate PBS for Europe-ancestry (CEU, TSI, GBR, and IBS, without FIN), we used YRI as an outgroup and East-Asian-ancestry (CHB, JPT, CHS, CDX, KHV) as a sister population; for East-Asian-ancestry, we used YRI as an outgroup and Europe-ancestry as a sister population; for YRI, we used East-Asian-ancestry as an outgroup and Europe-ancestry as a sister population.

### IL7 functional analyses

We PCR amplified and cloned the *IL7* wildtype (rs201412253-Val18) and mutant (rs201412253-18Ile) open reading frame (ORF) in the pcDNA5/FRT vector (ThermoFisher Scientific) using HindIII and BamHI restriction sites (see **Supplementary Table 22** for ORF and primer sequences). We validated the sequences of the two plasmids by Sanger Sequencing. Flip-In^TM^-293 cells (ThermoFisher Scientific) at 80% confluency were transfected with 1:10 mixes of empty pcDNA5 or pcDNA5 derivatives coding for IL7-Val8 or IL7-18Ile and pOG44 FLP recombinase coding vector (ThermoFisher Scientific) using polyethylenimine. Transfectant clones were expanded and selected in DMEM medium supplemented with 10% Foetal Bovine Serum, 4 mM L-glutamine, 100 IU penicillin, 100 μg/ml streptomycin and 100 μg/ml hygromycin. We measured the secretion of IL7 in eight independent clones for each *IL7* allele (rs201412253-Val18 and rs201412253-18Ile) as well as in four clones generated with the empty vector by ELISA assay. We used the High Sensitivity Quantikine HS ELISA kit from R & D Systems (Cat # HS750). We seeded 100,000 cells per 12-wells plates and grew them for 6 days in DMEM glutamax plus 10% FBS before doing the ELISA. We measured each supernatant in duplicate and seeded each of the clones in triplicate. The whole experiment was done on three different weeks (three complete biological replicates). We extracted total proteins from cells with RIPA buffer and we quantified the lysates by BCA. We used this quantification to normalize the ELISA assays. We extracted total RNA from ~500,000 cells using the Qiagen RNEasy kit (cat # 74136). We checked the quality of the RNA by Bioanalyzer and quantified its concentration by Nanodrop. We reverse transcribed 1 ug of total RNA into cDNA using the ABI kit (Life Technologies Cat # 4368814). We used two pairs of primers for *IL7* and assays for three normalizing genes (*HPRT*, *GAPDH*, *TBP,* **Supplementary Table 22**). We followed the MIQE recommendations and performed the qPCR reactions with the Sybergreen Platinum (Life Technologies Cat # 11733-046) on a Biorad CFX384 thermocycler.

## DATA AVAILABILITY STATEMENT

All results are available at the following URL: http://www.mhi-humangenetics.org/en/resources

## URLs

checkVCF: http://genome.sph.umich.edu/wiki/CheckVCF.py

EPACTS: https://genome.sph.umich.edu/wiki/EPACTS

Imputation servers: https://imputationserver.sph.umich.edu or https://imputation.sanger.ac.uk/

NCBI dbSNP: https://www.ncbi.nlm.nih.gov/projects/SNP/

PheKB: https://phekb.org/

Strand alignment resources: http://www.well.ox.ac.uk/~wrayner/strand

UK Biobank SAIGE results: http://pheweb.sph.umich.edu/SAIGE-UKB/

Variant Effect Predictor: https://useast.ensembl.org/info/docs/tools/vep/index.html

## References

Astle, W.J., Elding, H., Jiang, T., Allen, D., Ruklisa, D., Mann, A.L., Mead, D., Bouman, H., Riveros-Mckay, F., Kostadima, M.A., et al. (2016). The Allelic Landscape of Human Blood Cell Trait Variation and Links to Common Complex Disease. Cell 167, 1415–1429 e1419.

Auer, P.L., Johnsen, J.M., Johnson, A.D., Logsdon, B.A., Lange, L.A., Nalls, M.A., Zhang, G., Franceschini, N., Fox, K., Lange, E.M., et al. (2012). Imputation of exome sequence variants into population- based samples and blood-cell-trait-associated loci in African Americans: NHLBI GO Exome Sequencing Project. Am J Hum Genet 91, 794–808.

Auer, P.L., Teumer, A., Schick, U., O’Shaughnessy, A., Lo, K.S., Chami, N., Carlson, C., de Denus, S., Dube, M.P., Haessler, J., et al. (2014). Rare and low-frequency coding variants in CXCR2 and other genes are associated with hematological traits. Nat Genet.

Beutler, E., and West, C. (2005). Hematologic differences between African-Americans and whites: the roles of iron deficiency and alpha-thalassemia on hemoglobin levels and mean corpuscular volume. Blood 106, 740–745.

Brown, B.C., Asian Genetic Epidemiology Network Type 2 Diabetes, C., Ye, C.J., Price, A.L., and Zaitlen, N. (2016). Transethnic Genetic-Correlation Estimates from Summary Statistics. Am J Hum Genet 99, 76–88.

Bulik-Sullivan, B., Finucane, H.K., Anttila, V., Gusev, A., Day, F.R., Loh, P.R., ReproGen, C., Psychiatric Genomics, C., Genetic Consortium for Anorexia Nervosa of the Wellcome Trust Case Control, C., Duncan, L., et al. (2015a). An atlas of genetic correlations across human diseases and traits. Nat Genet 47, 1236–1241.

Bulik-Sullivan, B.K., Loh, P.R., Finucane, H.K., Ripke, S., Yang, J., Schizophrenia Working Group of the Psychiatric Genomics, C., Patterson, N., Daly, M.J., Price, A.L., and Neale, B.M. (2015b). LD Score regression distinguishes confounding from polygenicity in genome-wide association studies. Nat Genet 47, 291–295.

Bycroft, C., Freeman, C., Petkova, D., Band, G., Elliott, L.T., Sharp, K., Motyer, A., Vukcevic, D., Delaneau, O., O’Connell, J., et al. (2018). The UK Biobank resource with deep phenotyping and genomic data. Nature 562, 203–209.

Byrnes, J.R., and Wolberg, A.S. (2017). Red blood cells in thrombosis. Blood 130, 1795–1799.

Carroll, R.J., Bastarache, L., and Denny, J.C. (2014). R PheWAS: data analysis and plotting tools for phenome-wide association studies in the R environment. Bioinformatics 30, 2375–2376.

Chami, N., Chen, M.H., Slater, A.J., Eicher, J.D., Evangelou, E., Tajuddin, S.M., Love-Gregory, L., Kacprowski, T., Schick, U.M., Nomura, A., et al. (2016). Exome Genotyping Identifies Pleiotropic Variants Associated with Red Blood Cell Traits. Am J Hum Genet 99, 8–21.

Chen, X., Wang, H., Zhou, G., Zhang, X., Dong, X., Zhi, L., Jin, L., and He, F. (2009). Molecular population genetics of human CYP3A locus: signatures of positive selection and implications for evolutionary environmental medicine. Environ Health Perspect 117, 1541–1548.

Chen, Y., Fang, F., Hu, Y., Liu, Q., Bu, D., Tan, M., Wu, L., and Zhu, P. (2016). The Polymorphisms in LNK Gene Correlated to the Clinical Type of Myeloproliferative Neoplasms. PLoS One 11, e0154183.

Chu, S.G., Becker, R.C., Berger, P.B., Bhatt, D.L., Eikelboom, J.W., Konkle, B., Mohler, E.R., Reilly, M.P., and Berger, J.S. (2010). Mean platelet volume as a predictor of cardiovascular risk: a systematic review and meta-analysis. J Thromb Haemost 8, 148–156.

Colin, Y., Le Van Kim, C., and El Nemer, W. (2014). Red cell adhesion in human diseases. Current opinion in hematology 21, 186–192.

Corces, M.R., Buenrostro, J.D., Wu, B., Greenside, P.G., Chan, S.M., Koenig, J.L., Snyder, M.P., Pritchard, J.K., Kundaje, A., Greenleaf, W.J., et al. (2016). Lineage-specific and single-cell chromatin accessibility charts human hematopoiesis and leukemia evolution. Nat Genet 48, 1193–1203.

Das, S., Forer, L., Schonherr, S., Sidore, C., Locke, A.E., Kwong, A., Vrieze, S.I., Chew, E.Y., Levy, S., McGue, M., et al. (2016). Next-generation genotype imputation service and methods. Nat Genet 48, 1284–1287.

Delaneau, O., Zagury, J.F., and Marchini, J. (2013). Improved whole-chromosome phasing for disease and population genetic studies. Nat Methods 10, 5–6.

Denny, J.C., Ritchie, M.D., Basford, M.A., Pulley, J.M., Bastarache, L., Brown-Gentry, K., Wang, D., Masys, D.R., Roden, D.M., and Crawford, D.C. (2010). PheWAS: demonstrating the feasibility of a phenome-wide scan to discover gene-disease associations. Bioinformatics 26, 1205–1210.

Ding, K., de Andrade, M., Manolio, T.A., Crawford, D.C., Rasmussen-Torvik, L.J., Ritchie, M.D., Denny, J.C., Masys, D.R., Jouni, H., Pachecho, J.A., et al. (2013). Genetic variants that confer resistance to malaria are associated with red blood cell traits in African-Americans: an electronic medical record-based genome-wide association study. G3 (Bethesda) 3, 1061–1068.

Eicher, J.D., Chami, N., Kacprowski, T., Nomura, A., Chen, M.H., Yanek, L.R., Tajuddin, S.M., Schick, U.M., Slater, A.J., Pankratz, N., et al. (2016). Platelet-Related Variants Identified by Exomechip Meta-analysis in 157,293 Individuals. Am J Hum Genet 99, 40–55.

Evans, D.M., Frazer, I.H., and Martin, N.G. (1999). Genetic and environmental causes of variation in basal levels of blood cells. Twin Res 2, 250–257.

Fang, H., Hui, Q., Lynch, J., Honerlaw, J., Assimes, T.L., Huang, J., Vujkovic, M., Damrauer, S.M., Pyarajan, S., Gaziano, J.M., et al. (2019). Harmonizing Genetic Ancestry and Self-identified Race/Ethnicity in Genome-wide Association Studies. Am J Hum Genet 105, 763–772.

Gaziano, J.M., Concato, J., Brophy, M., Fiore, L., Pyarajan, S., Breeling, J., Whitbourne, S., Deen, J., Shannon, C., Humphries, D., et al. (2016). Million Veteran Program: A mega-biobank to study genetic influences on health and disease. J Clin Epidemiol 70, 214–223.

Genomes Project, C., Abecasis, G.R., Auton, A., Brooks, L.D., DePristo, M.A., Durbin, R.M., Handsaker, R.E., Kang, H.M., Marth, G.T., and McVean, G.A. (2012). An integrated map of genetic variation from 1,092 human genomes. Nature 491, 56–65.

Grinde, K.E., Qi, Q., Thornton, T.A., Liu, S., Shadyab, A.H., Chan, K.H.K., Reiner, A.P., and Sofer, T. (2019). Generalizing polygenic risk scores from Europeans to Hispanics/Latinos. Genet Epidemiol 43, 50–62.

Hinckley, J.D., Abbott, D., Burns, T.L., Heiman, M., Shapiro, A.D., Wang, K., and Di Paola, J. (2013). Quantitative trait locus linkage analysis in a large Amish pedigree identifies novel candidate loci for erythrocyte traits. Mol Genet Genomic Med 1, 131–141.

Johnson, K.E., and Voight, B.F. (2018). Patterns of shared signatures of recent positive selection across human populations. Nat Ecol Evol 2, 713–720.

Justice, A.E., Karaderi, T., Highland, H.M., Young, K.L., Graff, M., Lu, Y., Turcot, V., Auer, P.L., Fine, R.S., Guo, X., et al. (2019). Protein-coding variants implicate novel genes related to lipid homeostasis contributing to body-fat distribution. Nat Genet 51, 452–469.

Kanai, M., Akiyama, M., Takahashi, A., Matoba, N., Momozawa, Y., Ikeda, M., Iwata, N., Ikegawa, S., Hirata, M., Matsuda, K., et al. (2018). Genetic analysis of quantitative traits in the Japanese population links cell types to complex human diseases. Nat Genet 50, 390–400.

Kang, H.M., Sul, J.H., Service, S.K., Zaitlen, N.A., Kong, S.Y., Freimer, N.B., Sabatti, C., and Eskin, E. (2010). Variance component model to account for sample structure in genome-wide association studies. Nat Genet 42, 348–354.

Kimura, H., Ishibashi, T., Uchida, T., Maruyama, Y., Friese, P., and Burstein, S.A. (1990). Interleukin 6 is a differentiation factor for human megakaryocytes in vitro. European journal of immunology 20, 1927–1931.

Klarin, D., Damrauer, S.M., Cho, K., Sun, Y.V., Teslovich, T.M., Honerlaw, J., Gagnon, D.R., DuVall, S.L., Li, J., Peloso, G.M., et al. (2018). Genetics of blood lipids among ~300,000 multi-ethnic participants of the Million Veteran Program. Nat Genet 50, 1514–1523.

Kunishima, S., Kobayashi, R., Itoh, T.J., Hamaguchi, M., and Saito, H. (2009). Mutation of the beta1-tubulin gene associated with congenital macrothrombocytopenia affecting microtubule assembly. Blood 113, 458–461.

Li, Y.R., and Keating, B.J. (2014). Trans-ethnic genome-wide association studies: advantages and challenges of mapping in diverse populations. Genome Med 6, 91.

Lin, J., Zhu, Z., Xiao, H., Wakefield, M.R., Ding, V.A., Bai, Q., and Fang, Y. (2017). The role of IL-7 in Immunity and Cancer. Anticancer Res 37, 963–967.

Lo, K.S., Wilson, J.G., Lange, L.A., Folsom, A.R., Galarneau, G., Ganesh, S.K., Grant, S.F., Keating, B.J., McCarroll, S.A., Mohler, E.R., 3rd, et al. (2011). Genetic association analysis highlights new loci that modulate hematological trait variation in Caucasians and African Americans. Hum Genet 129, 307–317.

Loh, P.R., Kichaev, G., Gazal, S., Schoech, A.P., and Price, A.L. (2018). Mixed-model association for biobank-scale datasets. Nat Genet 50, 906–908.

Loh, P.R., Palamara, P.F., and Price, A.L. (2016). Fast and accurate long-range phasing in a UK Biobank cohort. Nat Genet 48, 811–816.

Lorenzo, F.R., Huff, C., Myllymaki, M., Olenchock, B., Swierczek, S., Tashi, T., Gordeuk, V., Wuren, T., Ri-Li, G., McClain, D.A., et al. (2014). A genetic mechanism for Tibetan high-altitude adaptation. Nat Genet 46, 951–956.

Magi, R., Horikoshi, M., Sofer, T., Mahajan, A., Kitajima, H., Franceschini, N., McCarthy, M.I., Cogent-Kidney Consortium, T.D. G.C., and Morris, A.P. (2017). Trans-ethnic meta-regression of genome-wide association studies accounting for ancestry increases power for discovery and improves fine-mapping resolution. Hum Mol Genet 26, 3639–3650.

Magi, R., and Morris, A.P. (2010). GWAMA: software for genome-wide association meta-analysis. BMC Bioinformatics 11, 288.

Mahajan, A., Taliun, D., Thurner, M., Robertson, N.R., Torres, J.M., Rayner, N.W., Payne, A.J., Steinthorsdottir, V., Scott, R.A., Grarup, N., et al. (2018). Fine-mapping type 2 diabetes loci to single-variant resolution using high-density imputation and islet-specific epigenome maps. Nat Genet 50, 1505–1513.

Marouli, E., Graff, M., Medina-Gomez, C., Lo, K.S., Wood, A.R., Kjaer, T.R., Fine, R.S., Lu, Y., Schurmann, C., Highland, H.M., et al. (2017). Rare and low-frequency coding variants alter human adult height. Nature 542, 186–190.

Marquez-Luna, C., Loh, P.R., South Asian Type 2 Diabetes, C., Consortium, S.T.D., and Price, A.L. (2017). Multiethnic polygenic risk scores improve risk prediction in diverse populations. Genet Epidemiol 41, 811–823.

Martin, A.R., Kanai, M., Kamatani, Y., Okada, Y., Neale, B.M., and Daly, M.J. (2019). Clinical use of current polygenic risk scores may exacerbate health disparities. Nat Genet 51, 584–591.

McCarthy, S., Das, S., Kretzschmar, W., Delaneau, O., Wood, A.R., Teumer, A., Kang, H.M., Fuchsberger, C., Danecek, P., Sharp, K., et al. (2016). A reference panel of 64,976 haplotypes for genotype imputation. Nat Genet 48, 1279–1283.

Mousas, A., Ntritsos, G., Chen, M.H., Song, C., Huffman, J.E., Tzoulaki, I., Elliott, P., Psaty, B.M., Blood-Cell, C., Auer, P.L., et al. (2017). Rare coding variants pinpoint genes that control human hematological traits. PLoS Genet 13, e1006925.

Nagai, A., Hirata, M., Kamatani, Y., Muto, K., Matsuda, K., Kiyohara, Y., Ninomiya, T., Tamakoshi, A., Yamagata, Z., Mushiroda, T., et al. (2017). Overview of the BioBank Japan Project: Study design and profile. J Epidemiol 27, S2–S8.

Pers, T.H., Timshel, P., and Hirschhorn, J.N. (2015). SNPsnap: a Web-based tool for identification and annotation of matched SNPs. Bioinformatics 31, 418–420.

Popejoy, A.B., and Fullerton, S.M. (2016). Genomics is failing on diversity. Nature 538, 161–164.

Popejoy, A.B., Ritter, D.I., Crooks, K., Currey, E., Fullerton, S.M., Hindorff, L.A., Koenig, B., Ramos, E.M., Sorokin, E.P., Wand, H., et al. (2018). The clinical imperative for inclusivity: Race, ethnicity, and ancestry (REA) in genomics. Hum Mutat 39, 1713–1720.

Rabbolini, D.J., Morel-Kopp, M.C., Chen, Q., Gabrielli, S., Dunlop, L.C., Chew, L.P., Blair, N., Brighton, T.A., Singh, N., Ng, A.P., et al. (2017). Thrombocytopenia and CD34 expression is decoupled from alpha-granule deficiency with mutation of the first growth factor-independent 1B zinc finger. J Thromb Haemost 15, 2245–2258.

Raffield, L.M., Ulirsch, J.C., Naik, R.P., Lessard, S., Handsaker, R.E., Jain, D., Kang, H.M., Pankratz, N., Auer, P.L., Bao, E.L., et al. (2018). Common alpha-globin variants modify hematologic and other clinical phenotypes in sickle cell trait and disease. PLoS Genet 14, e1007293.

Raj, T., Kuchroo, M., Replogle, J.M., Raychaudhuri, S., Stranger, B.E., and de Jager, P.L. (2013). Common risk alleles for inflammatory diseases are targets of recent positive selection. Am J Hum Genet 92, 517–529.

Rana, S.R., Sekhsaria, S., and Castro, O.L. (1993). Hemoglobin S and C traits: contributing causes for decreased mean hematocrit in African-American children. Pediatrics 91, 800–802.

Rappoport, N., Simon, A.J., Amariglio, N., and Rechavi, G. (2019). The Duffy antigen receptor for chemokines, ACKR1,- ‘Jeanne DARC’ of benign neutropenia. Br J Haematol 184, 497–507.

Reich, D., Nalls, M.A., Kao, W.H., Akylbekova, E.L., Tandon, A., Patterson, N., Mullikin, J., Hsueh, W.C., Cheng, C.Y., Coresh, J., et al. (2009). Reduced neutrophil count in people of African descent is due to a regulatory variant in the Duffy antigen receptor for chemokines gene. PLoS Genet 5, e1000360.

Roden, D.M., Pulley, J.M., Basford, M.A., Bernard, G.R., Clayton, E.W., Balser, J.R., and Masys, D.R. (2008). Development of a large-scale de-identified DNA biobank to enable personalized medicine. Clin Pharmacol Ther 84, 362–369.

Schick, U.M., Jain, D., Hodonsky, C.J., Morrison, J.V., Davis, J.P., Brown, L., Sofer, T., Conomos, M.P., Schurmann, C., McHugh, C.P., et al. (2016). Genome-wide Association Study of Platelet Count Identifies Ancestry-Specific Loci in Hispanic/Latino Americans. Am J Hum Genet 98, 229–242.

Tajuddin, S.M., Schick, U.M., Eicher, J.D., Chami, N., Giri, A., Brody, J.A., Hill, W.D., Kacprowski, T., Li, J., Lyytikainen, L.P., et al. (2016). Large-Scale Exome-wide Association Analysis Identifies Loci for White Blood Cell Traits and Pleiotropy with Immune-Mediated Diseases. Am J Hum Genet 99, 22–39.

Ulirsch, J.C., Lareau, C.A., Bao, E.L., Ludwig, L.S., Guo, M.H., Benner, C., Satpathy, A.T., Kartha, V.K., Salem, R.M., Hirschhorn, J.N., et al. (2019). Interrogation of human hematopoiesis at single-cell and single-variant resolution. Nat Genet 51, 683–693.

van Dongen, J., Jansen, R., Smit, D., Hottenga, J.J., Mbarek, H., Willemsen, G., Kluft, C., Collaborators, A., Penninx, B.W., Ferreira, M.A., et al. (2014). The contribution of the functional IL6R polymorphism rs2228145, eQTLs and other genome-wide SNPs to the heritability of plasma sIL-6R levels. Behav Genet 44, 368–382.

Wakefield, J. (2007). A Bayesian measure of the probability of false discovery in genetic epidemiology studies. Am J Hum Genet 81, 208–227.

Wakefield, J. (2009). Bayes factors for genome-wide association studies: comparison with P-values. Genet Epidemiol 33, 79–86.

Weir, B.S., and Cockerham, C.C. (1984). Estimating F-Statistics for the Analysis of Population Structure. Evolution 38, 1358–1370.

Wellcome Trust Case Control, C., Maller, J.B., McVean, G., Byrnes, J., Vukcevic, D., Palin, K., Su, Z., Howson, J.M., Auton, A., Myers, S., et al. (2012). Bayesian refinement of association signals for 14 loci in 3 common diseases. Nat Genet 44, 1294–1301.

Willer, C.J., Li, Y., and Abecasis, G.R. (2010). METAL: Fast and efficient meta-analysis of genomewide association scans. Bioinformatics 26, 2190–2191.

Williams, A.L., Patterson, N., Glessner, J., Hakonarson, H., and Reich, D. (2012). Phasing of many thousands of genotyped samples. Am J Hum Genet 91, 238–251.

Winkler, T.W., Day, F.R., Croteau-Chonka, D.C., Wood, A.R., Locke, A.E., Magi, R., Ferreira, T., Fall, T., Graff, M., Justice, A.E., et al. (2014). Quality control and conduct of genome-wide association meta-analyses. Nature protocols 9, 1192–1212.

Wojcik, G.L., Graff, M., Nishimura, K.K., Tao, R., Haessler, J., Gignoux, C.R., Highland, H.M., Patel, Y.M., Sorokin, E.P., Avery, C.L., et al. (2019). Genetic analyses of diverse populations improves discovery for complex traits. Nature 570, 514–518.

Xiang, K., Ouzhuluobu, Peng, Y., Yang, Z., Zhang, X., Cui, C., Zhang, H., Li, M., Zhang, Y., Bianba, et al. (2013). Identification of a Tibetan-specific mutation in the hypoxic gene EGLN1 and its contribution to high-altitude adaptation. Mol Biol Evol 30, 1889–1898.

Yi, X., Liang, Y., Huerta-Sanchez, E., Jin, X., Cuo, Z.X., Pool, J.E., Xu, X., Jiang, H., Vinckenbosch, N., Korneliussen, T.S., et al. (2010). Sequencing of 50 human exomes reveals adaptation to high altitude. Science 329, 75–78.

Zhernakova, A., Elbers, C.C., Ferwerda, B., Romanos, J., Trynka, G., Dubois, P.C., de Kovel, C.G., Franke, L., Oosting, M., Barisani, D., et al. (2010). Evolutionary and functional analysis of celiac risk loci reveals SH2B3 as a protective factor against bacterial infection. Am J Hum Genet 86, 970–977.

Zhou, W., Nielsen, J.B., Fritsche, L.G., Dey, R., Gabrielsen, M.E., Wolford, B.N., LeFaive, J., VandeHaar, P., Gagliano, S.A., Gifford, A., et al. (2018). Efficiently controlling for case-control imbalance and sample relatedness in large-scale genetic association studies. Nat Genet 50, 1335–1341.

