## Supplementary Information for "Trans-ethnic and ancestry-specific blood-cell genetics in 746,667 individuals from 5 global populations"

### Supplementary Methods

#### Conditional analyses in the UK Biobank European-ancestry population to identify independent variants associated with blood-cell traits

Stepwise multiple linear regression aims to identify the parsimonious subset of variants which explain the significant associations identified by univariate GWAS. For each blood index, the set of genome-wide significant variants was partitioned into the largest number of blocks such that no pair of blocks are separated by fewer than 5Mb, and no block contains more than 2,500 variants. Blocks are generated independently for each blood cell index. For each block we identify a parsimonious list of variants explaining the signal in that block using a stepwise conditional linear regression protocol discussed below. Linear regression is performed using *fastLM* from the R package *RcppEigen,* and run with a significance threshold of $8.31 \times 10^{-9}$. The stepwise linear regression proceeds in two stages, addition and removal of variants into to model which upon convergence represents the parsimonious signal in the block. The model starts with no variants and begins as follows iterating repeatedly through the following stages:

- **Initialisation step:** From the list of all variants in the block, add the variant with the lowest P-value that is also below the significance threshold ($8.31 \times 10^{-9}$).
- **Dropping:** Study the P-values for all variants in the model, if any of these are above the significance threshold we iteratively prune and rebuild model starting with the variant with the highest P-value. Once a variant is pruned it is returned to the list of variants not currently in the parsimonious model and may rejoin at a later iteration.
- **Addition:** Test each variant * not currently in the block sequentially in the model, add the variant with the lowest P-value which is below the threshold. Any tested variants which have a P-value of higher than 0.01 are not tested again in future iterations. * Variants are not permitted to be tested in the model if they have a LD $r^2 > 0.9$ with any variant currently in the model.
- **Completion:** If the algorithm could neither add a variant into the model nor remove a variant from the model then we abort the iteration with the model at this stage representing the parsimonious model for this block.

Following identification of conditionally significant variants in each block, all conditionally significant variants within each chromosome are put into a single linear model and tested with the same multiple stepwise linear regression algorithm as that defined above. The resultant set is the 'conditionally significant' list of variants for the blood cell index.

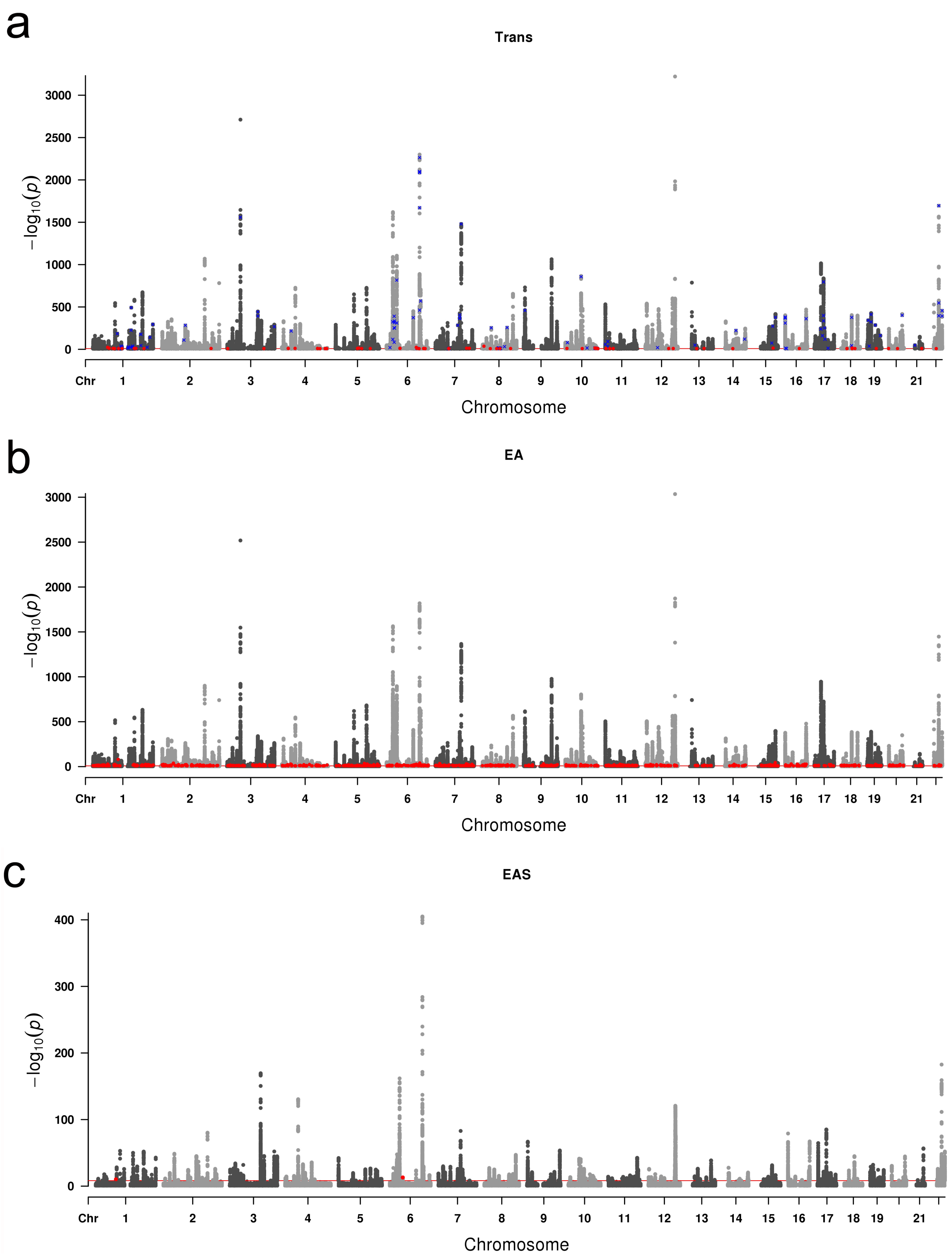
a
Trans
$-\log_{10}(p)$
3000
2500
2000
1500
1000
500
0
Chr 1 2 3 4 5 6 7 8 9 10 11 12 13 14 15 16 17 18 19 21
Chromosome
b
EA
$-\log_{10}(p)$
3000
2500
2000
1500
1000
500
0
Chr 1 2 3 4 5 6 7 8 9 10 11 12 13 14 15 16 17 18 19 21
Chromosome
c
EAS
$-\log_{10}(p)$
400
300
200
100
0
Chr 1 2 3 4 5 6 7 8 9 10 11 12 13 14 15 16 17 18 19 21
Chromosome

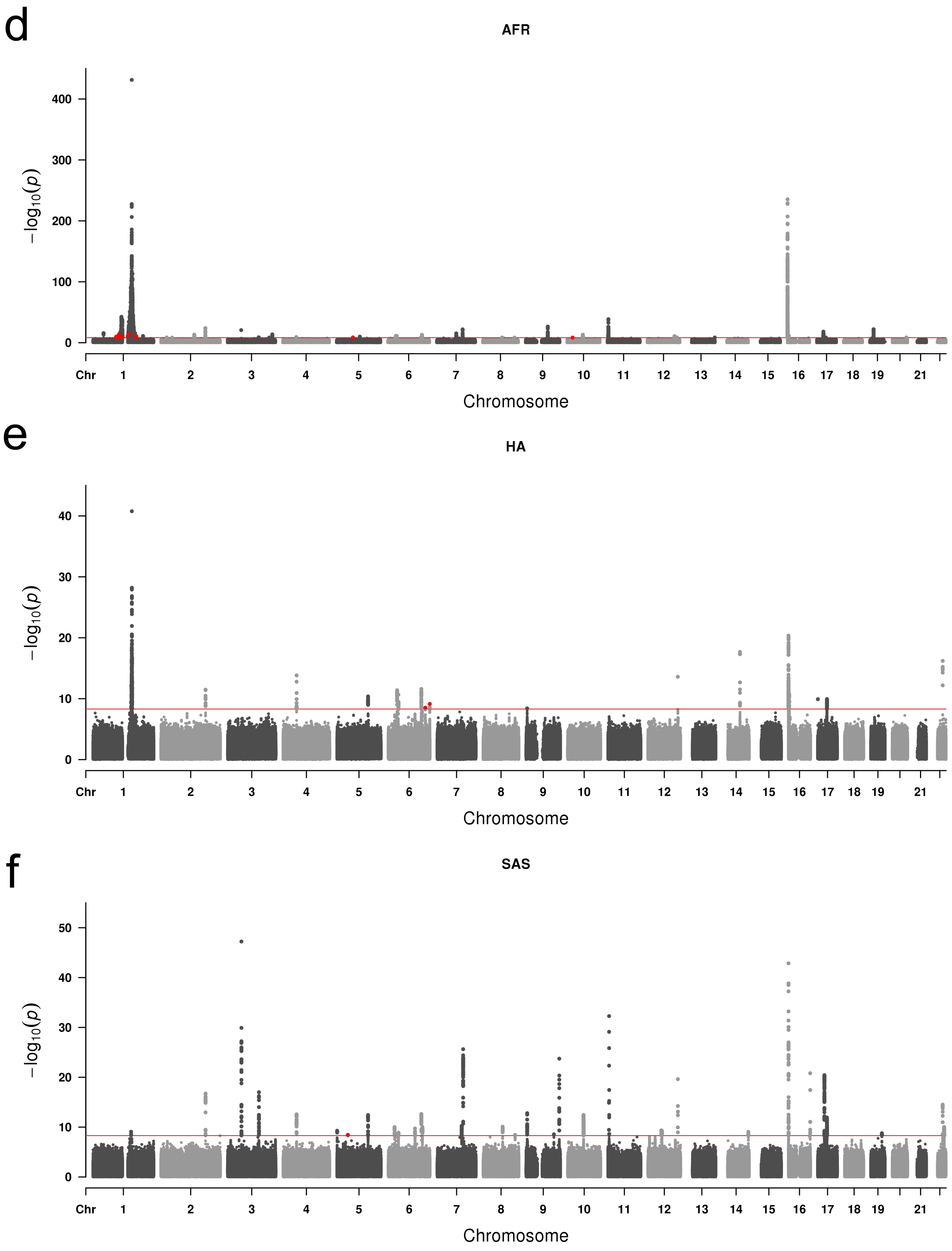


**Supplementary Figure 1.** Manhattan plots of association results in each population and in the trans-ethnic meta-analyses. At each variant, we report the smallest P-value across all 15 traits analyzed. In blue and red, we highlight novel loci and loci with heterogeneity P-value<$5x10^{-9}$

respectively. EA, European-ancestry; EAS, East-Asian-ancestry; HA, Hispanic-American-ancestry; AFR, African-ancestry; SAS, South-Asian ancestry.

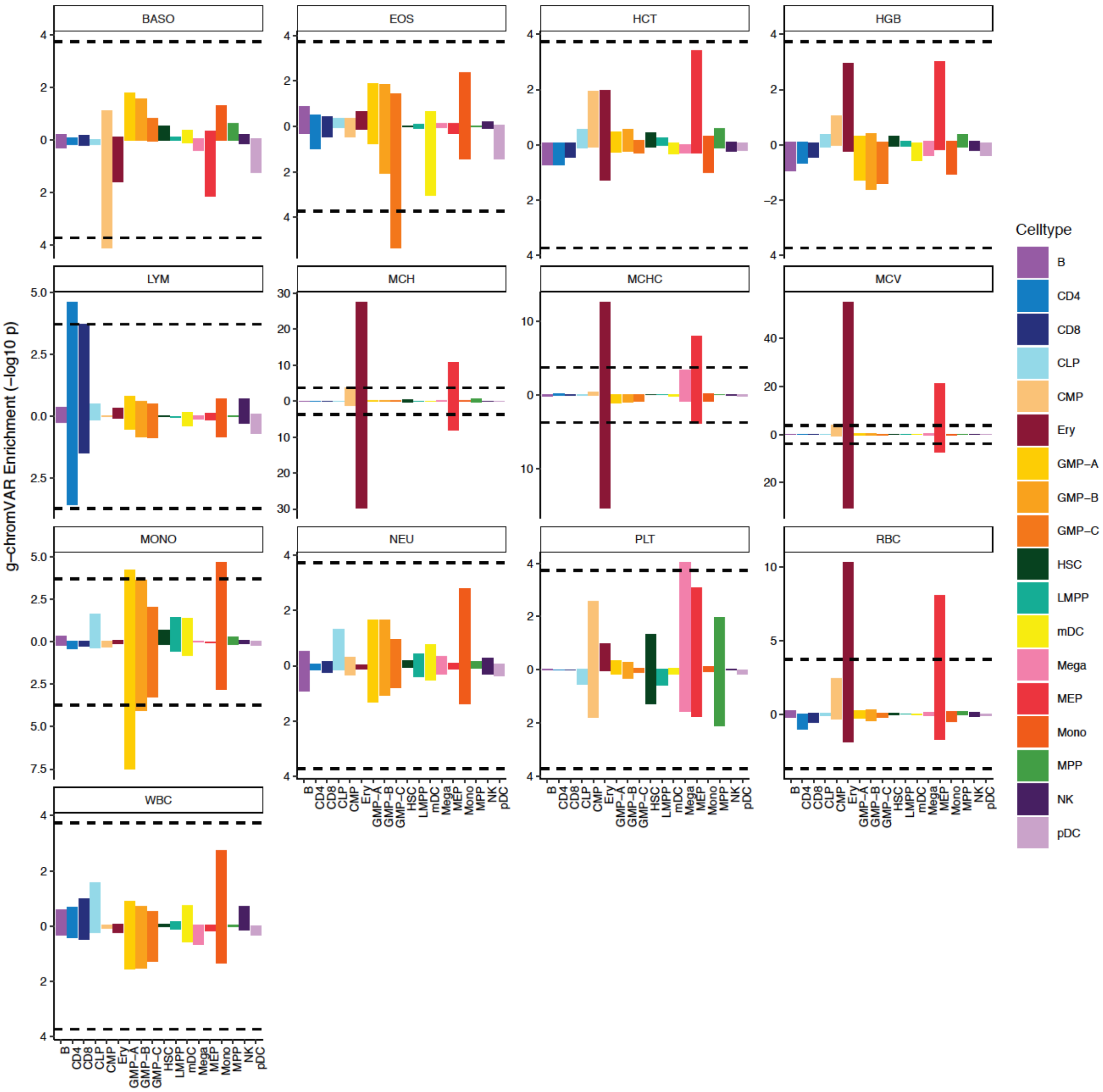


**Supplementary Figure 2.** Miami plot contrasting cell type enrichments of 95% credible set variants for 13 hematological traits studied in the European (top) and East Asian ancestry (bottom) GWAS, computed using g-chromVAR. The Bonferroni-adjusted significance level (one sided z-test) is indicated by the dotted line. mono, monocyte; gran, granulocyte; ery, erythroid; mega, megakaryocyte; CD4, $CD4^+$ T cell; CD8, $CD8^+$ T cell; B, B cell; NK, natural killer cell; mDC, myeloid dendritic cell; pDC, plasmacytoid dendritic cell; MPP, multipotent progenitor; LMPP, lymphoid-primed multipotent progenitor; CMP, common myeloid progenitor; CLP, common lymphoid progenitor; GMP, granulocyte–macrophage progenitor; MEP, megakaryocyte–erythroid progenitor.

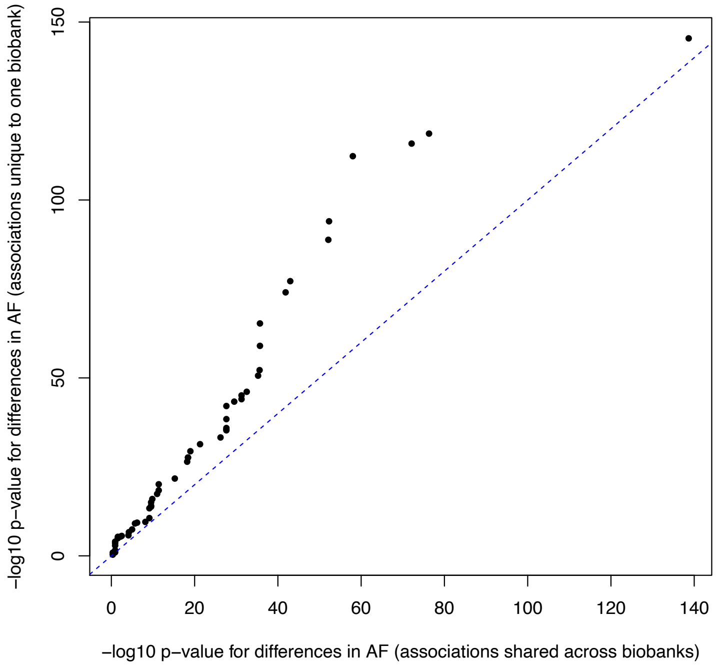


**Supplementary Figure 3.** Distribution of -$\log_{10}$(P-values) for testing differences in allele frequencies across populations. Quantiles of the distribution of variants with pheWAS effects shared across populations are plotted on the horizontal axis; the quantiles of the distribution of variants with pheWAS effects unique to a population are plotted on the vertical axis. AF, allele frequency.

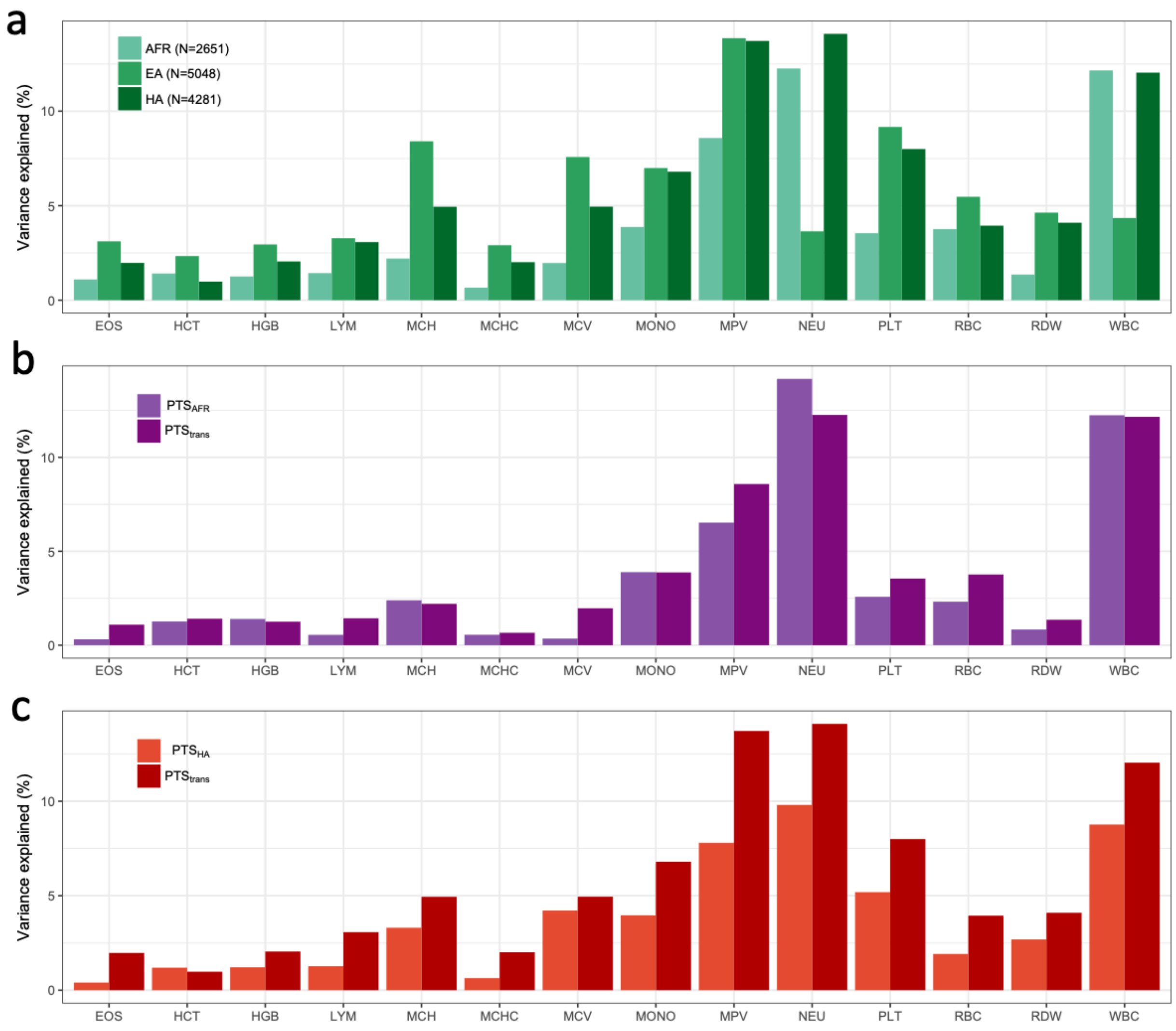


**Supplementary Figure 4.** Phenotypic variance explained by polygenic trait scores (PTS) in independent participants from the BioMe Biobank. Basophil counts were not tested as the PTS were not significant in any of the BioMe populations. (**a**) For each blood-cell trait, $PTS_{trans}$ were calculated using genome-wide significant variants identified in the trans-ethnic meta-analyses. Trait-increasing alleles were weighted using effect sizes derived from fixed-effect trans-ethnic meta-analyses. (**b**) In AFR BioMe participants, we compared the variance explained by $PTS_{trans}$ or $PTS_{AFR}$, a polygenic predictor calculated using the same trans-ethnic genome-wide significant variants, but weighted with AFR-specific effect sizes. (**c**) As for **b**, but for HA BioMe participants and using HA-specific effect sizes to weight $PTS_{HA}$.

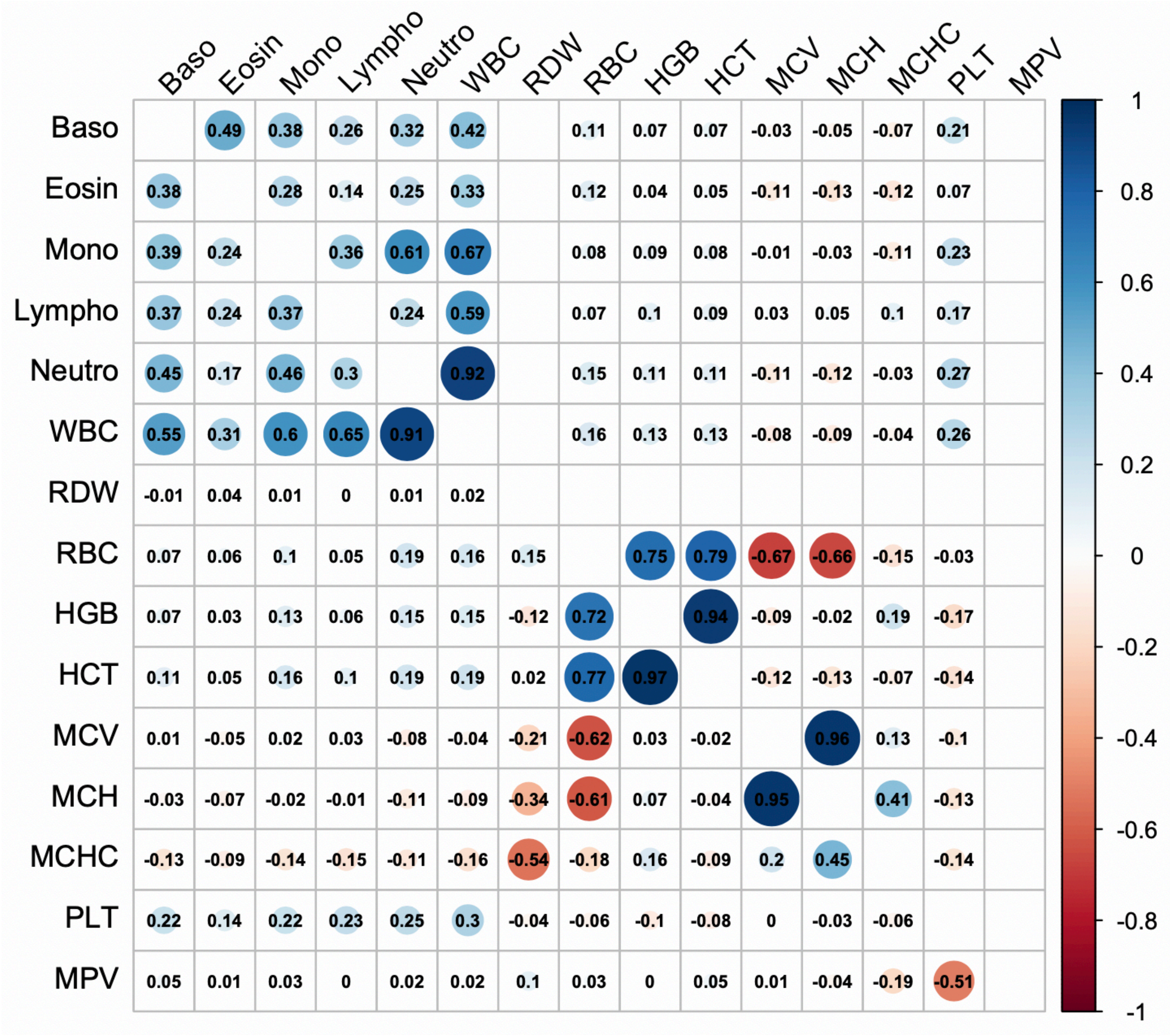

**Supplementary Figure 5.** Genetic correlations of blood-cell traits estimated using linkage disequilibrium score regression. The number in each cell correspond to the genetic correlation coefficient ($r_g$) between the pair of traits analyzed. Results over the diagonal (right side of the square) are for East Asians (EAS) whereas results under the diagonal (left side of the square) are for European-ancestry (EA) individuals. We note that the results on one side of the diagonal form an almost perfect mirror image of the results on the other side, indicating that the genetic correlations between pairs of blood-cell traits are very similar between EAS and EA. RDW and MPV results were not available in EAS.

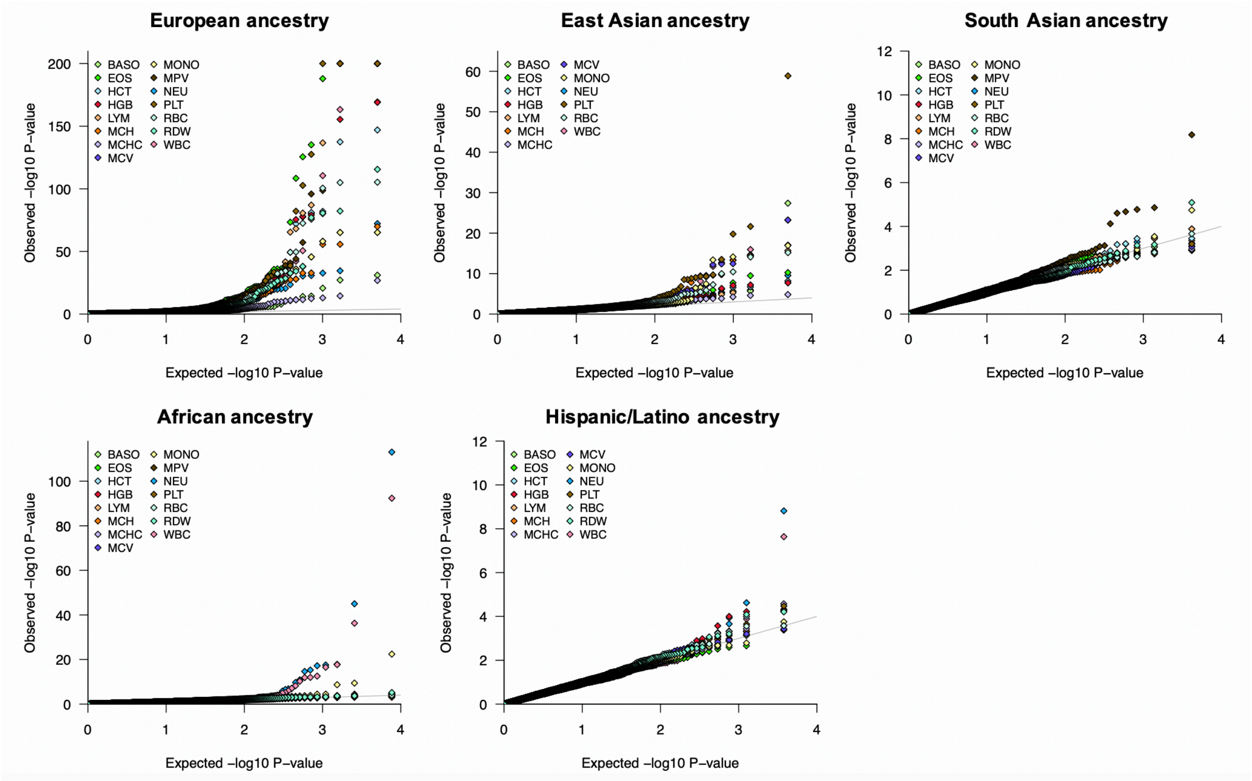


**Supplementary Figure 6.** Quantile-quantile plots of SNPs that "tag" selective sweeps in 1000 Genomes Project populations. For each of these variants, we retrieved the corresponding blood-cell trait association results from the ancestry-specific meta-analyses.

### VA Million Veteran Program membership

MVP Executive Committee

- Co-Chair: J. Michael Gaziano, M.D., M.P.H.
- Co-Chair: Rachel Ramoni, D.M.D., Sc.D.
- Jim Breeling, M.D. (ex-officio)
- Kyong-Mi Chang, M.D.
- Grant Huang, Ph.D.
- Sumitra Muralidhar, Ph.D.
- Christopher J. O'Donnell, M.D., M.P.H.
- Philip S. Tsao, Ph.D.

MVP Program Office

- Sumitra Muralidhar, Ph.D.
- Jennifer Moser, Ph.D.

MVP Recruitment/Enrollment

- Recruitment/Enrollment Director/Deputy Director, Boston – Stacey B. Whitbourne, Ph.D.; Jessica V. Brewer, M.P.H.
- MVP Coordinating Centers
    - Clinical Epidemiology Research Center (CERC), West Haven – John Concato, M.D., M.P.H.
    - Cooperative Studies Program Clinical Research Pharmacy Coordinating Center, Albuquerque - Stuart Warren, J.D., Pharm D.; Dean P. Argyres, M.S.
    - Genomics Coordinating Center, Palo Alto – Philip S. Tsao, Ph.D.
    - Massachusetts Veterans Epidemiology Research Information Center (MAVERIC), Boston - J. Michael Gaziano, M.D., M.P.H.
    - MVP Information Center, Canandaigua – Brady Stephens, M.S.
- Core Biorepository, Boston – Mary T. Brophy M.D., M.P.H.; Donald E. Humphries, Ph.D.
- MVP Informatics, Boston – Nhan Do, M.D.; Shahpoor Shayan
- Data Operations/Analytics, Boston – Xuan-Mai T. Nguyen, Ph.D.

MVP Science

- Genomics - Christopher J. O'Donnell, M.D., M.P.H.; Saiju Pyarajan Ph.D.; Philip S. Tsao, Ph.D.
- Phenomics - Kelly Cho, M.P.H, Ph.D.
- Data and Computational Sciences – Saiju Pyarajan, Ph.D.
- Statistical Genetics – Elizabeth Hauser, Ph.D.; Yan Sun, Ph.D.; Hongyu Zhao, Ph.D.

MVP Local Site Investigators

- Atlanta VA Medical Center (Peter Wilson)
- Bay Pines VA Healthcare System (Rachel McArdle)
- Birmingham VA Medical Center (Louis Dellitalia)

- Cincinnati VA Medical Center (John Harley)
- Clement J. Zablocki VA Medical Center (Jeffrey Whittle)
- Durham VA Medical Center (Jean Beckham)
- Edith Nourse Rogers Memorial Veterans Hospital (John Wells)
- Edward Hines, Jr. VA Medical Center (Salvador Gutierrez)
- Fayetteville VA Medical Center (Gretchen Gibson)
- VA Health Care Upstate New York (Laurence Kaminsky)
- New Mexico VA Health Care System (Gerardo Villareal)
- VA Boston Healthcare System (Scott Kinlay)
- VA Western New York Healthcare System (Junzhe Xu)
- Ralph H. Johnson VA Medical Center (Mark Hamner)
- Wm. Jennings Bryan Dorn VA Medical Center (Kathlyn Sue Haddock)
- VA North Texas Health Care System (Sujata Bhushan)
- Hampton VA Medical Center (Pran Iruvanti)
- Hunter Holmes McGuire VA Medical Center (Michael Godschalk)
- Iowa City VA Health Care System (Zuhair Ballas)
- Jack C. Montgomery VA Medical Center (Malcolm Buford)
- James A. Haley Veterans' Hospital (Stephen Mastorides)
- Louisville VA Medical Center (Jon Klein)
- Manchester VA Medical Center (Nora Ratcliffe)
- Miami VA Health Care System (Hermes Florez)
- Michael E. DeBakey VA Medical Center (Alan Swann)
- Minneapolis VA Health Care System (Maureen Murdoch)
- N. FL/S. GA Veterans Health System (Peruvemba Sriram)
- Northport VA Medical Center (Shing Shing Yeh)
- Overton Brooks VA Medical Center (Ronald Washburn)
- Philadelphia VA Medical Center (Darshana Jhala)
- Phoenix VA Health Care System (Samuel Aguayo)
- Portland VA Medical Center (David Cohen)
- Providence VA Medical Center (Satish Sharma)
- Richard Roudebush VA Medical Center (John Callaghan)
- Salem VA Medical Center (Kris Ann Oursler)
- San Francisco VA Health Care System (Mary Whooley)
- South Texas Veterans Health Care System (Sunil Ahuja)
- Southeast Louisiana Veterans Health Care System (Amparo Gutierrez)
- Southern Arizona VA Health Care System (Ronald Schifman)
- Sioux Falls VA Health Care System (Jennifer Greco)
- St. Louis VA Health Care System (Michael Rauchman)
- Syracuse VA Medical Center (Richard Servatius)
- VA Eastern Kansas Health Care System (Mary Oehlert)
- VA Greater Los Angeles Health Care System (Agnes Wallbom)
- VA Loma Linda Healthcare System (Ronald Fernando)
- VA Long Beach Healthcare System (Timothy Morgan)
- VA Maine Healthcare System (Todd Stapley)
- VA New York Harbor Healthcare System (Scott Sherman)
- VA Pacific Islands Health Care System (Gwenevere Anderson)

- VA Palo Alto Health Care System (Philip Tsao)
- VA Pittsburgh Health Care System (Elif Sonel)
- VA Puget Sound Health Care System (Edward Boyko)
- VA Salt Lake City Health Care System (Laurence Meyer)
- VA San Diego Healthcare System (Samir Gupta)
- VA Southern Nevada Healthcare System (Joseph Fayad)
- VA Tennessee Valley Healthcare System (Adriana Hung)
- Washington DC VA Medical Center (Jack Lichy)
- W.G. (Bill) Hefner VA Medical Center (Robin Hurley)
- White River Junction VA Medical Center (Brooks Robey)
- William S. Middleton Memorial Veterans Hospital (Robert Striker)

### Funding information and Conflicts of interests


| Author | Conflict of interest | Funding |
|---|---|---|
| Parsa Akbari | None | |
| Masato Akiyama | None | |
| William J. Astle | None | |
| Paul L. Auer | None | 1R01HL130733-01A1 |
| Erik L. Bao | None | ELB received support from the Howard Hughes Medical Institute Medical Research Fellowship |
| Traci M. Bartz | None | |
| Mélissa Beaudoin | None | |
| Yoav Ben-Shlomo | None | The Caerphilly Prospective Study was undertaken by the former MRC Epidemiology Unit (South Wales) and was funded by the Medical Research Council of the United Kingdom. The Caerphilly DNA bank was established by an MRC Grant (G9824960). |
| Andrew Beswick | None | |
| Jette Bork-Jensen | None | |
| Erwin P. Bottinger | None | |
| Jennifer A. Brody | None | |
| Linda Broer | None | The Rotterdam Study is funded by Erasmus Medical Center and Erasmus University, Rotterdam, Netherlands Organization for the Health Research and Development (ZonMw), the Research Institute for Diseases in the Elderly (RIDE), the Ministry of Education, Culture and Science, the Ministry for Health, Welfare and Sports, the European Commission (DG XII), and the Municipality of Rotterdam. The authors are grateful to the study participants, the staff from the Rotterdam Study and the participating general practitioners and pharmacists. |
| Adam S. Butterworth | ASB has received grants (outside of this work) from AstraZeneca, Biogen, BioMarin, Bioverativ, Merck, Novartis and Sanofi. | |

| | | |
|---|---|---|
| Cardiovascular Health Study (CHS) | None | This CHS research was supported by NHLBI contracts HHSN268201200036C, HHSN268200800007C, HHSN268201800001C, N01HC55222, N01HC85079, N01HC85080, N01HC85081, N01HC85082, N01HC85083, N01HC85086,R01HL068986; and NHLBI grants U01HL080295, R01HL087652, R01HL105756, R01HL103612, R01HL120393, and U01HL130114 with additional contribution from the National Institute of Neurological Disorders and Stroke (NINDS). Additional support was provided through R01AG023629 from the National Institute on Aging (NIA). A full list of principal CHS investigators and institutions can be found at CHS-NHLBI.org.<br>The provision of genotyping data was supported in part by the National Center for Advancing Translational Sciences, CTSI grant UL1TR001881, and the National Institute of Diabetes and Digestive and Kidney Disease Diabetes Research Center (DRC) grant DK063491 to the Southern California Diabetes Endocrinology Research Center.<br>The content is solely the responsibility of the authors and does not necessarily represent the official views of the National Institutes of Health. |
| Ming-Huei Chen | None | |
| Minhui Chen | None | |
| Charleston W.K. Chiang | None | |
| Kumaraswamynaidu Chitrala | None | |
| Kelly Cho | None | This research is based on data from the Million Veteran Program, Office of Research and Development, Veterans Health Administration, and was supported by awards MVP000 & I01-BX003340. |
| Hélène Choquet | None | Genotyping of the GERA cohort was funded by a grant from the National Institute on Aging, National Institute of Mental Health, and National Institute of Health Common Fund (RC2 AG036607). E.J. and H.C. are supported by National Eye Institute (NEI) grant R01 EY027004 (E.J.) and National Institute of Diabetes and Digestive and Kidney Diseases grant R01 DK116738 (E.J.) |
| Adolfo Correa | None | The Jackson Heart Study (JHS) is supported and conducted in collaboration with Jackson State University (HHSN268201800013I), Tougaloo College (HHSN268201800014I), the Mississippi State Department of Health (HHSN268201800015I) and the University of Mississippi Medical Center (HHSN268201800010I, HHSN268201800011I and HHSN268201800012I) contracts from the National Heart, Lung, and Blood Institute (NHLBI) and the National Institute on Minority Health and Health Disparities (NIMHD). The authors also wish to thank the staffs and participants of the JHS.<br>Disclaimer:<br>The following disclaimer must be included in your submitted manuscript: The views expressed in this manuscript are those of the authors and do not necessarily represent the views of the National Heart, Lung, and Blood Institute; the National Institutes of Health; or the U.S. Department of Health and Human Services. |
| John Danesh | None | J.D. is a British Heart Foundation Professor, European Research Council Senior Investigator, and National Institute for Health Research (NIHR) Senior Investigator. |
| Emanuele Di Angelantonio | None | |
| Niki Dimou | None | **Disclaimer:** Where authors are identified as personnel of the International Agency for Research on Cancer / World Health Organization, the authors alone are responsible for the views expressed in this article and they do not necessarily represent the decisions, policy or views of the International Agency for Research on Cancer / World Health Organization. |

| | | |
|---|---|---|
| Jingzhong Ding | None | |
| Paul Elliott | None | The Airwave Health Monitoring Study is funded by the UK Home Office, (grant number 780-TETRA) with additional support from the National Institute for Health Research Imperial College Biomedical Research Centre and UK Medical Research Council and Economic and Social Research Council (MR/S019669/1). We thank all participants in the Airwave Health Monitoring Study. This work used computing resources provided by the MRC- funded UK MEDical Bioinformatics partnership programme (UK MED-BIO) (MR/L01632X/1).<br>P.E. acknowledges support from the Medical Research Council (MRC) and Public Health England (PHE) Centre for Environment and Health (MR/L01341X/1 and MR/R023484/1); the NIHR Imperial College Biomedical Research Centre; and the NIHR Health Protection Research Unit in Health Impact of Environmental Hazards (HPRU-2012-10141). P.E. is a UK Dementia Research Institute (DRI) professor, at UK DRI at Imperial College London, funded by the MRC, Alzheimer's Society and Alzheimer's Research UK; and Associate Director of the Health Data Research UK (HDR-UK) London Centre funded by HDR UK Ltd sponsored by a consortium led by the UK Medical Research Council. |
| Tõnu Esko | None | |
| Evangelos Evangelou | None | |
| Michele K. Evans | None | |
| James S. Floyd | Has consulted for Shionogi Inc | This CHS research was supported by NHLBI contracts HHSN268201200036C, HSN268200800007C, HHSN268201800001C, N01HC55222, N01HC85079, N01HC85080, N01HC85081, N01HC85082, N01HC85083, N01HC85086; and NHLBI grants U01HL080295, R01HL085251, R01HL087652, R01HL105756, R01HL103612, R01HL120393, and U01HL130114 with additional contribution from the National Institute of Neurological Disorders and Stroke (NINDS). Additional support was provided through R01AG023629 from the National Institute on Aging (NIA). A full list of principal CHS investigators and institutions can be found at CHS-NHLBI.org. The provision of genotyping data was supported in part by the National Center for Advancing Translational Sciences, CTSI grant UL1TR001881, and the National Institute of Diabetes and Digestive and Kidney Disease Diabetes Research Center (DRC) grant DK063491 to the Southern California Diabetes Endocrinology Research Center. The content is solely the responsibility of the authors and does not necessarily represent the official views of the National Institutes of Health. |
| Jean-François Gauchat | None | |
| Mohsen Ghanbari | None | The Rotterdam Study is funded by Erasmus Medical Center and Erasmus University, Rotterdam, Netherlands Organization for the Health Research and Development (ZonMw), the Research Institute for Diseases in the Elderly (RIDE), the Ministry of Education, Culture and Science, the Ministry for Health, Welfare and Sports, the European Commission (DG XII), and the Municipality of Rotterdam. The authors are grateful to the study participants, the staff from the Rotterdam Study and the participating general practitioners and pharmacists. |
| Niels Grarup | None | |
| Andreas Greinacher | None | SHIP is part of the Community Medicine Research net of the University of Greifswald, Germany (www.community-medicine.de), which is funded by the Federal Ministry of Education and Research (grants no. 01ZZ9603, 01ZZ0103, and 01ZZ0403), the Siemens AG, the Ministry of Cultural Affairs as well as the Social Ministry of the Federal State of Mecklenburg-West Pomerania, and the network 'Greifswald Approach to Individualized Medicine (GANI_MED)' funded by the Federal Ministry of Education and Research (grant |

| | | |
|---|---|---|
| | | 03IS2061A). ExomeChip data have been supported by the Federal Ministry of Education and Research (grant no. 03Z1CN22) and the Federal State of Mecklenburg-West Pomerania. |
| Michael H. Guo | None | |
| Jeff Haessler | None | |
| Torben Hansen | None | |
| Joanna M. M. Howson | During the drafting of the manuscript, JMMH became a fulltime employee of Novo Nordisk | British Heart Foundation (RG/13/13/30194; RG/18/13/33946) and the National Institute for Health Research[Cambridge Biomedical Research Centre at the Cambridge University Hospitals NHS Foundation Trust] [*]. |
| Wei Huang | None | We thank the National Institute for Nutrition and Health, Chinese Center for Disease Control and Prevention, the Chinese National Human Genome Center at Shanghai, the Carolina Population Center, the University of North Carolina at Chapel Hill, and all of the participants and study investigators involved in the China Health and Nutrition Survey. Data collection and analysis was supported by the Carolina Population Center (P2C HD050924, T32 HD007168), the NIH (R01HD30880, R01 DK056350, R24 HD050924, R01 HD38700, R01 DK072193 and U01 DK105561), the NIH Fogarty International Center (D43 TW009077, D43 TW007709), the China Ministry of Health, the Chinese National Human Genome Center at Shanghai, the China-Japan Friendship Hospital, and the Beijing Municipal Center for Disease Prevention and Control. |
| Jennifer E. Huffman | None | This research is based on data from the Million Veteran Program, Office of Research and Development, Veterans Health Administration, and was supported by awards MVP000 & I01-BX003340. |
| Tao Jiang | None | |
| Andrew D. Johnson | None | This research was supported by NHLBI Intramural Research Funding to ADJ. The views expressed in this manuscript are those of the authors and do not necessarily represent the views of the National Heart, Lung, and Blood Institute, the NIH, or the U.S. Department of Health and Human Services. The NHLBI's Framingham Heart Study is a joint project of the National Institutes of Health and Boston University School of Medicine and was supported by contract N01-HC-25195. |
| Eric Jorgenson | None | Genotyping of the GERA cohort was funded by a grant from the National Institute on Aging, National Institute of Mental Health, and National Institute of Health Common Fund (RC2 AG036607). E.J. and H.C. are supported by National Eye Institute (NEI) grant R01 EY027004 (E.J.) and National Institute of Diabetes and Digestive and Kidney Diseases grant R01 DK116738 (E.J.) |
| Tim Kacprowski | None | SHIP is part of the Community Medicine Research net of the University of Greifswald, Germany (www.community-medicine.de), which is funded by the Federal Ministry of Education and Research (grants no. 01ZZ9603, 01ZZ0103, and 01ZZ0403), the Siemens AG, the Ministry of Cultural Affairs as well as the Social Ministry of the Federal State of Mecklenburg-West Pomerania, and the network 'Greifswald Approach to Individualized Medicine (GANI_MED)' funded by the Federal Ministry of Education and Research (grant |

| | | |
|---|---|---|
| | | 03IS2061A). ExomeChip data have been supported by the Federal Ministry of Education and Research (grant no. 03Z1CN22) and the Federal State of Mecklenburg-West Pomerania. |
| Mika Kähönen | None | The Finnish Cardiovascular Study (FINCAVAS) has been financially supported by the Competitive Research Funding of the Tampere University Hospital (Grant 9M048 and 9N035), the Finnish Cultural Foundation, the Finnish Foundation for Cardiovascular Research, the Emil Aaltonen Foundation, Finland, the Tampere Tuberculosis Foundation, EU Horizon 2020 (grant 755320 for TAXINOMISIS), and the Academy of Finland grant 322098. |
| Yoichiro Kamatani | None | |
| Masahiro Kanai | None | |
| Savita Karthikeyan | None | |
| Fotis Koskeridis | None | |
| Leslie A. Lange | None | |
| Véronique Laplante | None | |
| Caleb A. Lareau | None | |
| Terho Lehtimäki | None | The Young Finns Study has been financially supported by the Academy of Finland: grants 322098, 286284, 134309 (Eye), 126925, 121584, 124282, 129378 (Salve), 117787 (Gendi), and 41071 (Skidi); the Social Insurance Institution of Finland; Competitive State Research Financing of the Expert Responsibility area of Kuopio, Tampere and Turku University Hospitals (grant X51001); Juho Vainio Foundation; Paavo Nurmi Foundation; Finnish Foundation for Cardiovascular Research ; Finnish Cultural Foundation; The Sigrid Juselius Foundation; Tampere Tuberculosis Foundation; Emil Aaltonen Foundation; Yrjö Jahnsson Foundation; Signe and Ane Gyllenberg Foundation; Diabetes Research Foundation of Finnish Diabetes Association; EU Horizon 2020 (grant 755320 for TAXINOMISIS); European Research Council (grant 742927 for MULTIEPIGEN project); and Tampere University Hospital Supporting Foundation. |
| Markus M. Lerch | None | SHIP is part of the Community Medicine Research net of the University of Greifswald, Germany (www.community-medicine.de), which is funded by the Federal Ministry of Education and Research (grants no. 01ZZ9603, 01ZZ0103, and 01ZZ0403), the Siemens AG, the Ministry of Cultural Affairs as well as the Social Ministry of the Federal State of Mecklenburg-West Pomerania, and the network 'Greifswald Approach to Individualized Medicine (GANI_MED)' funded by the Federal Ministry of Education and Research (grant 03IS2061A). ExomeChip data have been supported by the Federal Ministry of Education and Research (grant no. 03Z1CN22) and the Federal State of Mecklenburg-West Pomerania. |
| Guillaume Lettre | None | We thank all participants and staff of the André and France Desmarais MHI Biobank. This work was funded by the Canadian Institutes of Health Research (PJT #156248), the Canada Research Chair Program, Genome Quebec and Genome Canada, and the Montreal Heart Institute Foundation. |
| Yun Li | None | R01 HL129132. Genotyping of the GERA cohort was funded by a grant from the National Institute on Aging, National Institute of Mental Health, and National Institute of Health Common Fund (RC2 AG036607). |
| Bingshan Li | None | U01HG009086 |
| Allan Linneberg | None | |

| | | |
|---|---|---|
| Yongmei Liu | None | |
| Ken Sin Lo | None | |
| Ruth J.F. Loos | None | Ruth Loos is supported by funds of the NIH (R01DK110113; R01DK107786; R01HL142302; R01DK110113) |
| Leo-Pekka Lyytikäinen | None | The Young Finns Study has been financially supported by the Academy of Finland: grants 322098, 286284, 134309 (Eye), 126925, 121584, 124282, 129378 (Salve), 117787 (Gendi), and 41071 (Skidi); the Social Insurance Institution of Finland; Competitive State Research Financing of the Expert Responsibility area of Kuopio, Tampere and Turku University Hospitals (grant X51001); Juho Vainio Foundation; Paavo Nurmi Foundation; Finnish Foundation for Cardiovascular Research ; Finnish Cultural Foundation; The Sigrid Juselius Foundation; Tampere Tuberculosis Foundation; Emil Aaltonen Foundation; Yrjö Jahnsson Foundation; Signe and Ane Gyllenberg Foundation; Diabetes Research Foundation of Finnish Diabetes Association; EU Horizon 2020 (grant 755320 for TAXINOMISIS); European Research Council (grant 742927 for MULTIEPIGEN project); and Tampere University Hospital Supporting Foundation. |
| Regina Manansala | None | 1R01HL130733-01A1 |
| Ani Manichaikul | None | Support for the statistical analyses in MESA was provided by R01 HL120393 and R01 HL105756. |
| Arden Mascoti | None | |
| Koichi Matsuda | None | |
| MESA/MESA SHARe | None | Support for the statistical analyses in MESA was provided by R01 HL120393 and R01 HL105756. MESA and the MESA SHARe project are conducted and supported by the National Heart, Lung, and Blood Institute (NHLBI) in collaboration with MESA investigators. Support for MESA is provided by contracts HHSN268201500003I, N01-HC-95159, N01-HC-95160, N01-HC-95161, N01-HC-95162, N01-HC-95163, N01-HC-95164, N01-HC-95165, N01-HC-95166, N01-HC-95167, N01-HC-95168, N01-HC-95169, UL1-TR-000040, UL1-TR-001079, UL1-TR-001420. The provision of genotyping data was supported in part by the National Center for Advancing Translational Sciences, CTSI grant UL1TR001881, and the National Institute of Diabetes and Digestive and Kidney Disease Diabetes Research Center (DRC) grant DK063491 to the Southern California Diabetes Endocrinology Research Center. Funding for SHARe genotyping was provided by NHLBI Contract N02-HL-64278. Genotyping was performed at Affymetrix (Santa Clara, California, USA) and the Broad Institute of Harvard and MIT (Boston, Massachusetts, USA) using the Affymetrix Genome-Wide Human SNP Array 6.0. |
| Karen L. Mohlke | None | |
| Nina Mononen | None | The Finnish Cardiovascular Study (FINCAVAS) has been financially supported by the Competitive Research Funding of the Tampere University Hospital (Grant 9M048 and 9N035), the Finnish Cultural Foundation, the Finnish Foundation for Cardiovascular Research, the Emil Aaltonen Foundation, Finland, the Tampere Tuberculosis Foundation, EU Horizon 2020 (grant 755320 for TAXINOMISIS), and the Academy of Finland grant 322098. |
| Abdou Mousas | None | |
| Yoshinori Murakami | None | |
| Girish N. Nadkarni | None | |
| Matthias Nauck | None | SHIP is part of the Community Medicine Research net of the University of Greifswald, Germany (www.community-medicine.de), which is funded by the |

| | | |
|---|---|---|
| | | Federal Ministry of Education and Research (grants no. 01ZZ9603, 01ZZ0103, and 01ZZ0403), the Siemens AG, the Ministry of Cultural Affairs as well as the Social Ministry of the Federal State of Mecklenburg-West Pomerania, and the network 'Greifswald Approach to Individualized Medicine (GANI_MED)' funded by the Federal Ministry of Education and Research (grant 03IS2061A). ExomeChip data have been supported by the Federal Ministry of Education and Research (grant no. 03Z1CN22) and the Federal State of Mecklenburg-West Pomerania. |
| Kjell Nikus | None | The Finnish Cardiovascular Study (FINCAVAS) has been financially supported by the Competitive Research Funding of the Tampere University Hospital (Grant 9M048 and 9N035), the Finnish Cultural Foundation, the Finnish Foundation for Cardiovascular Research, the Emil Aaltonen Foundation, Finland, the Tampere Tuberculosis Foundation, EU Horizon 2020 (grant 755320 for TAXINOMISIS), and the Academy of Finland grant 322098. |
| Yukinori Okada | None | |
| Willem H. Ouwehand | None | W.H.O. is a NIHR Senior Investigator |
| Nathan Pankratz | None | |
| Oluf Pedersen | None | |
| Michael Preuss | None | |
| Bruce M. Psaty | None | |
| Huijun Qian | None | |
| Laura M. Raffield | None | T32 HL129982 |
| Olli T. Raitakari | None | The Young Finns Study has been financially supported by the Academy of Finland: grants 322098, 286284, 134309 (Eye), 126925, 121584, 124282, 129378 (Salve), 117787 (Gendi), and 41071 (Skidi); the Social Insurance Institution of Finland; Competitive State Research Financing of the Expert Responsibility area of Kuopio, Tampere and Turku University Hospitals (grant X51001); Juho Vainio Foundation; Paavo Nurmi Foundation; Finnish Foundation for Cardiovascular Research ; Finnish Cultural Foundation; The Sigrid Juselius Foundation; Tampere Tuberculosis Foundation; Emil Aaltonen Foundation; Yrjö Jahnsson Foundation; Signe and Ane Gyllenberg Foundation; Diabetes Research Foundation of Finnish Diabetes Association; EU Horizon 2020 (grant 755320 for TAXINOMISIS); European Research Council (grant 742927 for MULTIEPIGEN project); and Tampere University Hospital Supporting Foundation. |
| Alexander P. Reiner | None | APR is supported by the National Heart, Lung, and Blood Institute, National Institutes of Health (R01HL129132 and R01HL130733). The WHI program is funded by the National Heart, Lung, and Blood Institute, National Institutes of Health, U.S. Department of Health and Human Services through contracts HHSN268201600018C, HHSN268201600001C, HHSN268201600002C, HHSN268201600003C, and HHSN268201600004C. For a list of all the investigators who have contributed to WHI science, please visit: https://www.whi.org/researchers/Documents%20%20Write%20a%20Paper/WHI%20Investigator%20Long%20List.pdf |
| Stephen S. Rich | None | |
| David J. Roberts | None | D.J.R. was supported by the NIHR Programme "Erythropoiesis in Health and Disease" (NIHR-RP-PG-0310-1004). |

| | | |
|---|---|---|
| Benjamin A.T. Rodriguez | None | |
| Jonathan D. Rosen | None | |
| Jerome I. Rotter | None | |
| Saori Sakaue | None | |
| Vijay G. Sankaran | None | |
| Petra Schubert | None | This research is based on data from the Million Veteran Program, Office of Research and Development, Veterans Health Administration, and was supported by awards MVP000 & I01-BX003340. |
| Nicole Soranzo | None | NS is supported by the Wellcome Trust, the BHF and the NIHR |
| Cassandra N. Spracklen | None | CNS was supported by American Heart Association Postdoctoral Fellowship 15POST24470131 and 17POST33650016 |
| Praveen Surendran | None | |
| Hua Tang | None | |
| Jean-Claude Tardif | None | |
| Frank J.A. van Rooij | None | The Rotterdam Study is funded by Erasmus Medical Center and Erasmus University, Rotterdam, Netherlands Organization for the Health Research and Development (ZonMw), the Research Institute for Diseases in the Elderly (RIDE), the Ministry of Education, Culture and Science, the Ministry for Health, Welfare and Sports, the European Commission (DG XII), and the Municipality of Rotterdam. The authors are grateful to the study participants, the staff from the Rotterdam Study and the participating general practitioners and pharmacists. |
| Uwe Völker | None | SHIP is part of the Community Medicine Research net of the University of Greifswald, Germany (www.community-medicine.de), which is funded by the Federal Ministry of Education and Research (grants no. 01ZZ9603, 01ZZ0103, and 01ZZ0403), the Siemens AG, the Ministry of Cultural Affairs as well as the Social Ministry of the Federal State of Mecklenburg-West Pomerania, and the network 'Greifswald Approach to Individualized Medicine (GANI_MED)' funded by the Federal Ministry of Education and Research (grant 03IS2061A). ExomeChip data have been supported by the Federal Ministry of Education and Research (grant no. 03Z1CN22) and the Federal State of Mecklenburg-West Pomerania. |
| Henry Völzke | None | SHIP is part of the Community Medicine Research net of the University of Greifswald, Germany (www.community-medicine.de), which is funded by the Federal Ministry of Education and Research (grants no. 01ZZ9603, 01ZZ0103, and 01ZZ0403), the Siemens AG, the Ministry of Cultural Affairs as well as the Social Ministry of the Federal State of Mecklenburg-West Pomerania, and the network 'Greifswald Approach to Individualized Medicine (GANI_MED)' funded by the Federal Ministry of Education and Research (grant 03IS2061A). ExomeChip data have been supported by the Federal Ministry of Education and Research (grant no. 03Z1CN22) and the Federal State of Mecklenburg-West Pomerania. |
| Dragana Vuckovic | None | |
| Nicholas A. Watkins | None | |

| | | |
|---|---|---|
| Peter W.F. Wilson | None | This research is based on data from the Million Veteran Program, Office of Research and Development, Veterans Health Administration, and was supported by awards MVP000 & I01-BX003340. |
| Xue Zhong | None | |
| Alan B. Zonderman | None | |
